# Collagen Tubular Airway-on-Chip for Extended Epithelial Culture and Investigation of Ventilation Dynamics

**DOI:** 10.1101/2023.10.05.561125

**Authors:** Wuyang Gao, Kayshani R. Kanagarajah, Emma Graham, Kayla Soon, Teodor Veres, Theo J. Moraes, Christine E. Bear, Ruud A. Veldhuizen, Amy P. Wong, Axel Günther

## Abstract

The lower respiratory tract is a hierarchical network of compliant tubular structures that are made from extracellular matrix proteins with a wall lined by an epithelium. While microfluidic airway-on-a-chip models incorporate the effects of shear and stretch on the epithelium, week-long air-liquid-interface (ALI) culture remains limited to static conditions. The circular cross-section and substrate compliance associated with intact airways have yet to be recapitulated to allow studies of epithelial injuries under physiological and ventilation conditions. To overcome these limitations, we present a collagen tube-based airway model. Sustaining a functional human bronchial epithelium during two-week perfusion is accomplished by continuously supplying warm, humid air at the apical side and culture medium at the basal side. The model faithfully recapitulates human airways in size, composition, and mechanical microenvironment, allowing for the first time dynamic studies of elastocapillary phenomena associated with regular breathing as well as mechanical ventilation, along with the impact on epithelial cells. Findings reveal the epithelium to become increasingly damaged when subjected to repetitive collapse and reopening as opposed to overdistension and suggest expiratory flow resistance to reduce atelectasis. We expect the model to find broad potential applications in organ-on-a-chip applications for various tubular tissues.

## 1. Introduction

The human lung contains a hierarchical network of tubular airways with lumens lined by resident pulmonary epithelial cells that are continuously exposed to a complex biomechanical environment, including wall shear stress (**Figure 1A**), compression, stretch, and even collapse of the airway wall (**Figure 1B**).^[1]^ The disruption of the physiological conditions affects the epithelium in different ways that include apoptosis,^[2]^ proliferation,^[3]^ surfactant dysfunction,^[4,5]^ inflammation,^[6]^ impaired mucociliary transport,^[7]^ which may lead to fibrosis^[8]^ and affect the pathogenesis and progression of many pulmonary diseases such as acute respiratory distress syndrome (ARDS) and ventilator induced lung injury (VILI).^[1,9]^

**Figure 1.**
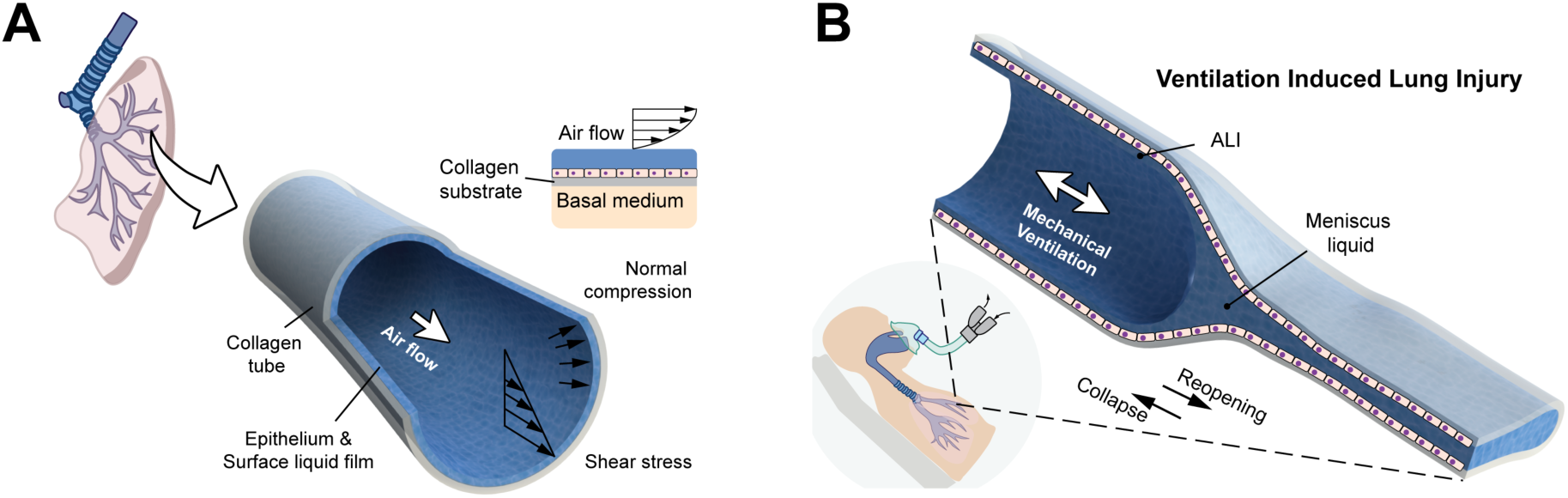
Schematic representation of lower airway biomechanical microenvironment during normal breathing and mechanical ventilation. (A) Schematic of airway microenvironment in native lung. (B) Schematic illustrating reopening of collapsed airway by propagating air plug during ventilation induced lung injury (VILI).

During the past 40 years,^[10]^ airway epithelial cells have been routinely cultured on synthetic membrane surfaces under static air-liquid interface (ALI) conditions, allowing maturation and differentiation during long-term culture,^[11,12]^ apicobasal polarization along with the investigation of barrier function,^[13]^ mucociliary dynamics,^[14–16]^ and air compression effect.^[9,17]^ Airway-on-a-chip models introduced during the past 15 years^[18,19]^ utilize microfluidic networks, in many cases separated by a rigid synthetic membrane, to present the epithelium with air and culture medium at the respective sides. These in vitro models emulate the local structure (e.g., hydraulic diameter), respiratory motion (e.g., cyclic stretch), immune cell recruitment, and spatial organization of epithelial cells with other cell types (detailed summary provided in **Table S1, Supporting Information**). However, the epithelium obtained on the apical side of these models is planar, unlike the curvature of the airway lumen. While week-long culture of epithelia has been demonstrated, it is so far largely limited to static air conditions, as the introduction of physiological airflow resulted in both cell viability and function to decline within only a few days.^[18,20,21]^

Notably, in the context of VILI, elevated lung volume may result in localized overdistension (“volutrauma”), leading to alveolar rupture, air leaks, and gross barotrauma. Conversely, during periods of low lung volume, injuries of the epithelium are caused by the repetitive collapse and reopening of small airways or alveolar ducts (“atelectrauma”).^[22]^ These phenomena are influenced by different parameters during mechanical ventilation, including the positive end-expiratory pressure (PEEP),^[23]^ the tidal volume,^[24,25]^ and the expiratory flow resistance.^[26]^ Collapse is often observed at the end of expiration, followed by reopening at the beginning of the next inspiration. Ex vivo experiments performed on isolated murine^[27]^ and canine^[28]^ lung lobes suggest two mechanisms for this phenomenon: “meniscus formation” and “compliant” collapse and reopening (**Figure 1B**).^[29]^ Previous in vitro studies investigated air plug propagation in parallel-plate flow chambers to simulate injuries related to the first mechanism. Slower air plug velocities^[30]^ and stiffer substrates^[31]^ were found to increase the injury of the epithelium, while application of surfactant mitigated the effect. The Takayama group^[19,32,33]^ investigated the injury of a planar epithelium using an airway-on-a-chip device that was supported by a synthetic membrane while a liquid plug confined between two air plugs propagated over it. Studies related to the second mechanism have not yet been feasible as airway-on-a-chip devices so far rely on polydimethylsiloxane (PDMS), plastic, or other stiff substrate materials (i.e., the substrate elastic modulus exceeded 1 MPa^[34,35]^) to define airway channel walls – several orders of magnitude too stiff to recapitulate airway collapse dynamics during mechanical ventilation.

Here, we report an advanced airway-on-a-chip model to overcome these limitations. Utilizing the previously reported extrusion of collagen-based tubular structures,^[36]^ we present: (1) A collagen tube based airway model for the human bronchial and alveolar ducts embedded with a custom designed microfluidic device; (2) The capacity of tuning the compliance and elasticity of the airway model wall within physiologically relevant ranges by tailoring collagen source and incubation protocol; (3) Confluent human bronchial epithelia lining the lumen of the tubular airway model, which developed and maintained under physiological air flow for up to 14 days; (4) Three ALI conditions: static, unidirectional, and bidirectional airflow, allow the investigation of airway dynamics during mechanical ventilation including shear stress, transmural pressure, tidal volume, cyclic stretch, effect of surfactant, expiratory flow resistance, as well as repetitive collapse and reopening; (5) The unique collapsible, stretchable and biocompatible nature of our tubular airway model allows combined studies of ventilation dynamics and resulting tissue injury.

## 2. Results and Discussion

### 2.1. Microfluidic Collagen Tube Hosting Device

**Figure 2A** shows a tube segment that was prepared from 10 mg ml^-1^ type I bovine hide collagen solution as previously described.^[36]^ We designed and fabricated a multi-layer thermoplastic hosting device, manually sutured the collagen tube segment onto it (**Figure 2B**). This configuration enables independent perfusion through tube lumen and superfusion from the outside with individual flowrates (for air or culture medium) and small transmural pressures (with a resolution of approximately 2 mmH_2_O), as depicted in **Figure 2C**. Two reservoirs were incorporated to seed cellsat a high yield to the inside of the tube, and to maintain the osmotic pressure in the culture medium during long-term ALI culture, respectively (detailed explanations are provided in **Section S2** and **Section S6, Supporting Information**). **Figures 2D** and **2E** show assembled and exploded views of the tube hosting device, fabricated from five individual poly (methyl methacrylate) (PMMA) layers. Microfluidic channel networks within the respective layers were patterned by laser cutting, followed by thermal bonding between all layers in the presented order. Luer lock adapters and syringe filters were attached to external medium/air perfusion and ensured sterility. Further information on the design, fabrication, optical access, and sterilization processes is summarized in **Section S2**, **Supporting Information**.

**Figure 2.**
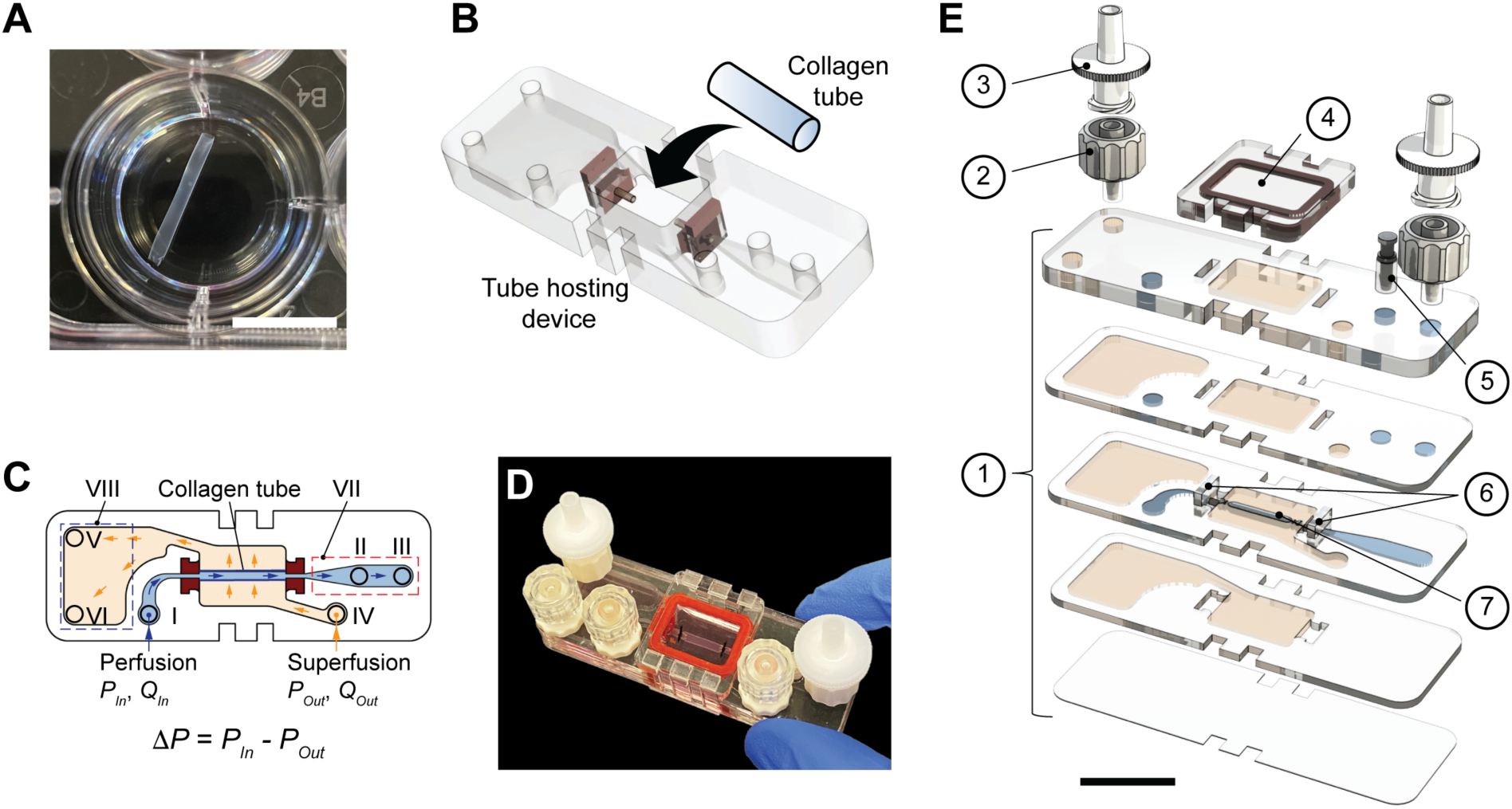
Airway-on-chip hosting device and perfusion platform. (A) Extruded collagen tube manually dissected into approximately 15 mm-long segment (inner diameter 1.7 mm, thickness 27 (m). Scale bar: 10 mm. (B) Schematic of collagen tube cannulated to thermoplastic hosting device. (C) Schematic of independent perfusion (➔) and superfusion (➔) within tube hosting device. (I ∼ III) perfusion portals, (IV ∼ VI) superfusion portals, (VII) epithelial cell seeding reservoir, (VIII) superfusion reservoir for maintaining osmotic pressure during long-term ALI culture. Transmural pressure across collagen tube-based airway model set by difference between perfusion and superfusion pressure. (D) Photograph of assembled airway-on-chip. (E) Exploded view of 5-layer thermoplastic tube hosting device including ① thermoplastic layers patterned with custom designed microchannel networks, ② Luer lock adapters connecting perfusion and superfusion, ③ syringe filter with 0.22 μm pore size, ④ reversible lid with molded PDMS O-ring, ⑤ plug for cell seeding reservoir, ⑥ canula adapters (Figure S1F, Supporting Information) containing polytetrafluoroethylene (PTFE) dispensing needles to connect collagen tube segment ⑦. Scale bar: 20 mm.

### 2.2. Mechanical Strength and Compliance of Collagen Tube

After cannulating to the hosting device, we applied a previously described incubation protocol^[37,38]^ to further enhance the mechanical strength of the tubular structure. As summarized in **Table 1** and **Figure 3**, four types of tubes were prepared by varying the collagen source and the incubation protocol after extrusion. At identical conditions (pH = 2 and collagen concentration 10 mg ml^-1^), we found that tubes prepared from collagen I derived from bovine hide (sample II) exhibited 0.4 times thinner walls, compared with the ones prepared from rat tail collagen I (sample I). When comparing samples II and I, Young’s moduli were 25 times and 7 times higher in the axial and circumferential directions, respectively. We attribute the this difference to the discrepancies in the viscosity and gelation kinetics between the two collagen solutions during tube formation. Briefly, during co-axial extrusion, the slower gelation of bovine collagen extended the period of time during which water was continuously osmotically removed from collagen solution via macromolecular crowding. Consequently, thinner walled tubes were formed and higher collagen fiber densities obtained.^[36]^ Samples I and II exhibited strain-to-failure ratios exceeding 70%, while displaying a burst pressure below 4 kPa. Further air-drying (sample III) and crosslinking with genipin (sample IV) enhanced the stiffness of the chip hosted collagen tube segment, exhibiting a burst pressure up to 26.1 ± 12.0 kPa. The presented data demonstrate the capacity to vary mechanical properties of chip hosted collagen tube segments over several orders of magnitude (e.g., Young’s moduli varied from 50 kPa to 15 MPa). This tunability is desirable when selecting tubular structures with specific stiffnesses and compliance for the target tissue or organs. Considering tube strength, elasticity, permeability, and optical access (avoiding autofluorescence), the remainder of our work is focused on tubes prepared from bovine collagen (referred to as sample II).

**Figure 3.**
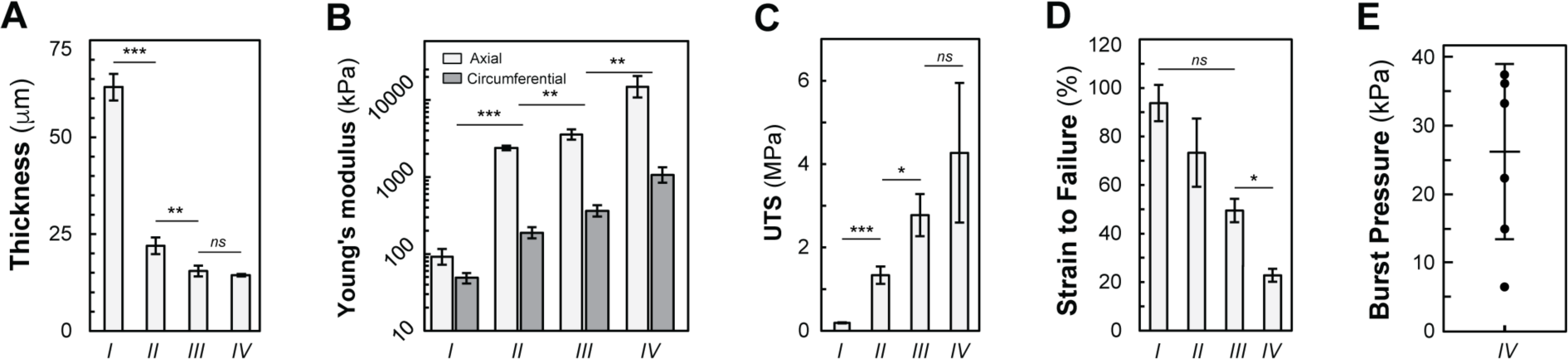
Tunable mechanical properties of chip-hosted collagen tubes. (A) Tube wall thickness, (B) Young’s moduli in axial and circumferential directions, (C) Ultimate tensile strength (UTS) in axial direction, (D) Strain to failure, and (E) burst pressure for tube samples prepared according to conditions in Table 1. Data presented as mean ± standard deviation, N = 4-6. ***p<0.001, ** p<0.01, * p<0.05.

**Table 1.**
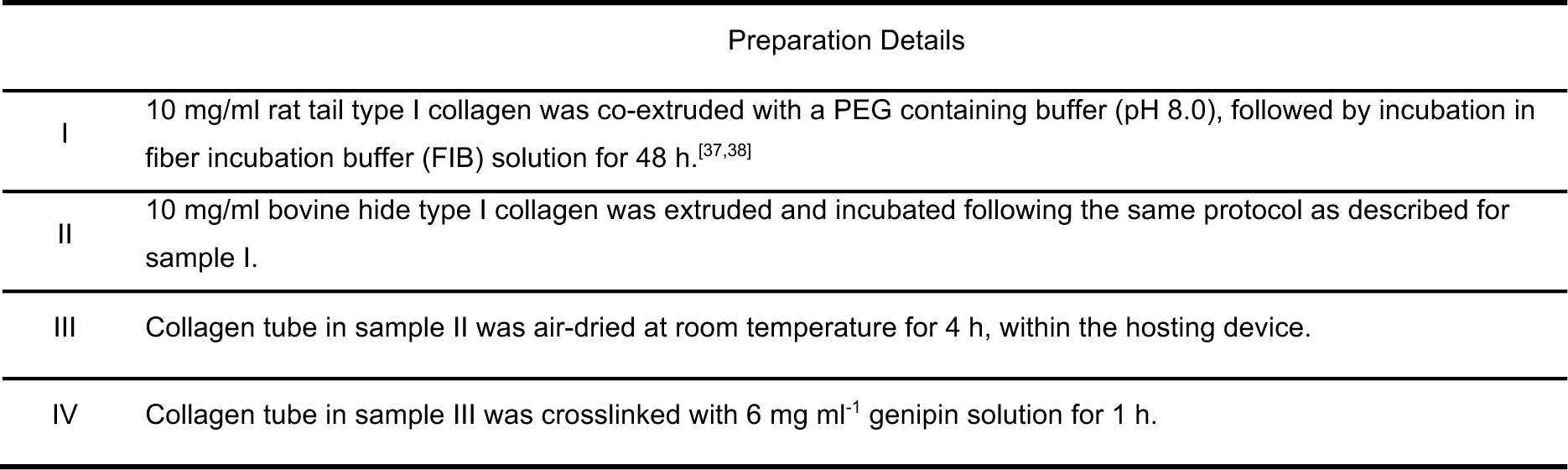
Collagen tube sample preparation information.

### 2.3. Bronchial Epithelium Culture at ALI under Physiological Airflow

**Figure 4A** summarizes the wall shear stress distribution at different branching orders within the human respiratory system, adapted from Weibel.^[39]^ **Figure 4B** provides a summary of the microfluidic airway models reported in the literature (including both synthetic^[18,19,32]^ or protein-based substrates^[40,41]^), with emphasis on culture duration and airflow conditions. While many efforts reported short-term (typically less than 2 days) ALI culture durations, the present work achieves week-long dynamic ALI culture of our collagen tube-based airway model. First, we used a numerical simulation with a commercial finite element solver (ANSYS, Canonsburg, PA, USA, Release 18.1) to investigate the impact of buoyancy on the soft collagen tube configuration and the variation of strain along its axis (**Section S4**, **Supporting Information**). To achieve a uniform strain distribution (coefficient of variation < 2.5%) at ι1P < 50 mmH_2_O, collagen tube lengths ranging from 10 to 12 mm were selected for the experiments reported in this paper (**Figure S3D**, **Supporting Information**).

**Figure 4.**
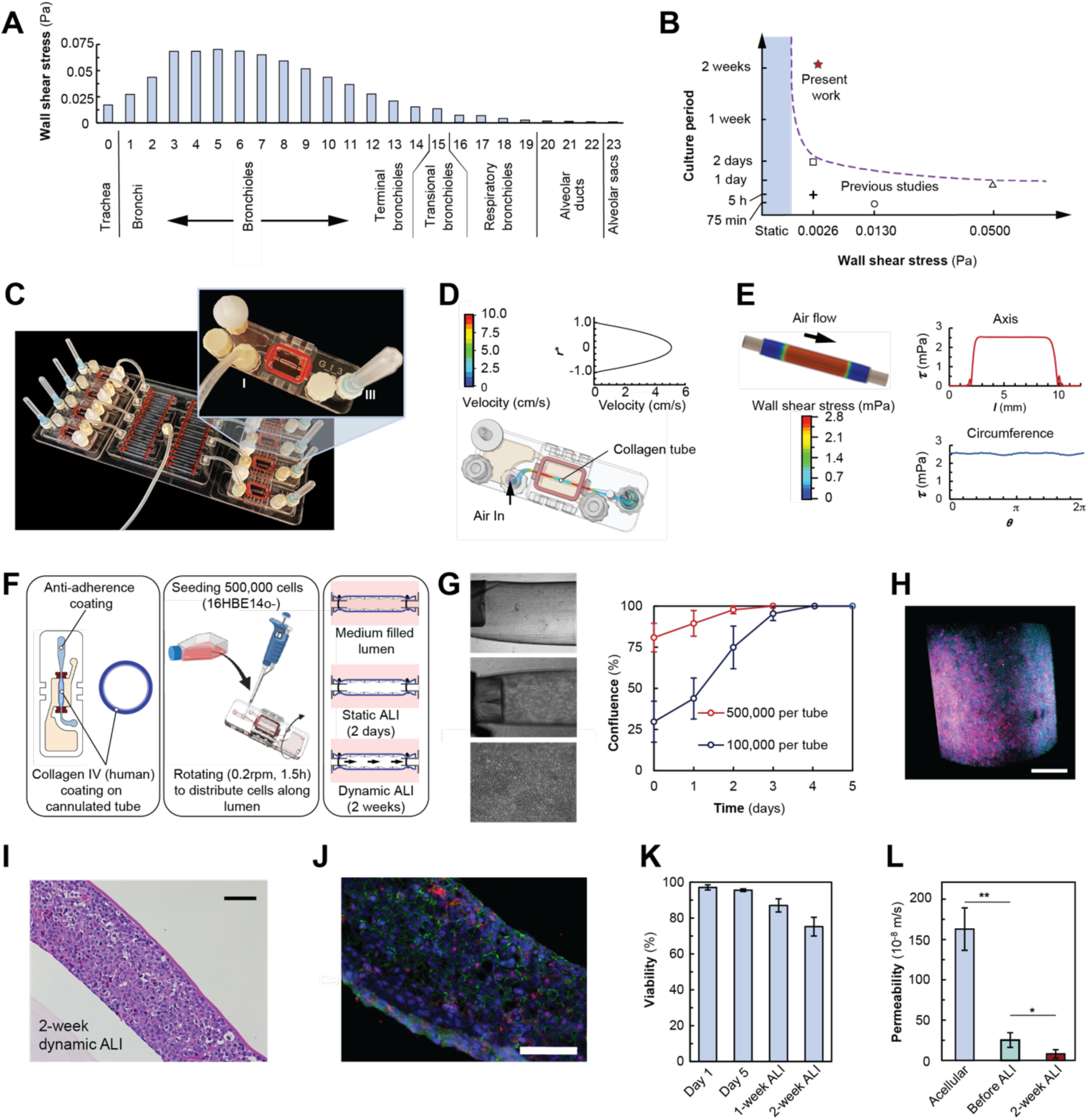
Long term dynamic ALI culture in collagen tube based airway-on-chip model. (A) Wall shear stress for different generations of airways in human respiratory system. (B) Applied wall shear stress and duration of ALI culture reported in different airway-on-chip platforms (□^[41]^, +^[19]^, ○^[44]^, △^[20]^) comparted with present work (*). (C) Photograph of platform accommodating eight collagen-based airway-on-chip devices during dynamic ALI culture. (D) Numerically predicted streamlines and velocity profiles during air perfusion of collagen tube (average velocity 2.5 cm/s). (E) Simulated distribution of wall shear stress along axis (-) and circumference (-). (F) Schematic illustration of experimental sequence (**Section S6**, **Supporting Information** for detail) involving cell seeding into chip-hosted and collagen IV coated collagen tube, rotation to promote uniform cell attachment, followed by culture with liquid-filled lumen, and static and dynamic ALI. (G) Bright-field micrographs on left correspond to acellular tube (4×, top) and cellular tube under confluent conditions (4×, center, and 20×, bottom). Increasing confluency during 5 days of culture shown on right. (H) 3D image of Pan-cytokeratin (magenta) and nucleus (blue) for HBE cells on lumen. Scale bar: 500 μm. (I) H&E staining of dissected cellular collagen tube after 2-week dynamic ALI culture. (J) Immunostaining of epithelium after 2-week ALI culture with unidirectionally applied 2.5 cm/s average air velocity. Nucleus (blue), ZO-1 (green), Ki67 (red). Scale bar: 75 μm. (L) Viability from cell seeding to 2-week culture. (M) Permeability of acellular (blue) and cellular collagen tube with confluent HBE cells prior to (green) and after 2-week ALI culture (red). Graphs represent mean ± standard error, N = 3-4. ** p<0.01, * p<0.05.

**Figure 4C** and **Section S3** (**Supporting Information**) illustrate the air perfusion through the tube lumen, where air enters the hosting device from port (I) and exits from port (III). At a designed flowrate of 3.6 ml min^-1^, **Figure 4D** shows streamlines of the velocity field within the tube lumen, as well as the velocity distribution in the radial direction, corresponding to an average air velocity of 2.5 cm/s. **Figure 4E** shows the corresponding shear stress distribution along the length of the tube segment. Except for the regions proximal to the two suture locations, uniform shear distribution of approximately 0.026 mPa was achieved for two thirds of the length. Small non-uniformities along the circumference were caused by small deviations from a circular cross-section due to buoyancy. Computed wall shear stresses closely match those observed in the airways of generations 19-20 in the adult human respiratory system during relaxed breathing (**Figure 4A**). Unlike current rectangular-channel airway-on-a-chip models, our model distributes the wall shear stress uniformly along the circumference, closely resembling native airways. (**Figure S4**, **Supporting Information**). The corresponding tube lumen pressure distribution is shown in **Figure S2B** (**Supporting Information**). To avoid the hosted collagen tube from collapsing due due to capillary pressure during air perfusion, a 1-inch long, 21-gauge needle was attached downstream of port (III) to increase the luminal pressure (and thereby the transmural pressure) by approximately 92 Pa. Accurate control of small transmural pressure levels is essential, as transmural pressures exceeding 300 Pa were already found to gradually increase the collagen tube diameter (Tube sample II) over time and ultimately result in tube rupture within only a few days of ALI culture.

To establish physiologically relevant and consistent dynamic ALI culture conditions on the luminal surface of multiple tubes, a custom-designed air-flow system was constructed and provided control over the air flowrate, the transmural pressure, the temperature, and the humidity, as shown in **Figure S2C** (**Supporting Information**). The air supplied to multiple tube hosting devices was conditioned and regulated using a series of in-line particle filters, a pressure controller, an air humidifier, a heater, a separator for condensate, as well as flow resistors positioned upstream of individual tube hosting devices. These components ensured the delivery of sterile, humid (relative humidity > 95%) and warm (36.5-37.2°C) air. The air-flow system was designed to accommodate and perfuse 32 single tube hosting devices in parallel (**Figure 4C** and **Figure S2C**, **Supporting Information**).

To demonstrate the capacity of our airway-on-chip model to support week-long ALI culture, a human bronchial epithelium was established on the luminal surface of chip hosted collagen tube segment. **Figure 4F** illustrates the protocol that includes the following steps: (1) Fabrication and sterilization of tube hosting device. To achieve high cell seeding yield, the reservoir (VII) was initially rinsed with an anti-adherence solution to prevent undesired cell adhesion in this specific area. (2) Manual cannulation of collagen tube segments, filling the lumen with type IV collagen-derived from human placenta^[11]^ and depositing a uniform coating by controlled air finger propagation through the lumen (for detailed protocol see **Figure S6**, **Supporting Information**) ultilizing a classical coating phenomenon^[42]^ instead of a previously demonstrated viscous fingering effect.^[43]^ (3) Seeding and culture of cells in culture medium-filled lumen until confluency. Either 100,000 or 500,000 human bronchial epithelial (HBE) cells were then seeded to one tube and distributed circumferentially by rotating the hosting device around the tube axis (detailed cell seeding procedure described in **Figure S7B**, **Supporting Information**). (4) Transition to static ALI and dynamic 2 week-ALI culture.

As shown in **Figure 4G**, normal proliferation of HBE cells was observed on the lumen and a confluent epithelium obtained 4 - 5 days after seeding and culture in medium filled lumen. 3D Light sheet confocal imaging (**Figure 4H**) revealed integrated epithelia. After obtaining confluency, static ALI culture was performed for two days, followed by 2-week dynamic ALI culture with warm, humid air. During the latter, the medium within the superfusion reservoir was manually exchanged every other day. Importantly, distilled water was added daily to maintain a constant osmotic pressure (**Figure S8B-E**, **Supporting Information**). **Figure 4I** and **4J** show H&E and immunostaining images of bronchial epithelia after 2-week unidirectional air perfusion with an average (bulk) velocity of 2.5 cm s^-1^. More than =70% of cells remained viable from day 1 after cell seeding until after the two-week dynamic ALI culture (**Figure 4K**). Barrier function of the epithelium was assessed by supplying the fluorescent marker fluorescein isothiocyanate-dextran (4 kDa FITC dextran flux assay, **Section S7**, **Supporting Information**) to the lumen. Due to permeation across the epithelium and tube wall, the FITC dextran intensity increases at the superfusion side (**Figure S9B**, **Supporting Information**). Fluorescent time-lapse images of this increase were recorded and converted to the permeability values shown in **Figure 4L**. The observed reduction in the permeability over time is attributed to the progressive maturation of the epithelial barrier during the two-week dynamic ALI culture.

### 2.4. Dynamics of ALI

Experiments were performed for three ALI conditions: static air, unidirectional airflow, and bidirectional airflow. As shown in **Figure 5A** and **5B**, the collapse and reopening at static ALI conditions was studied by continuously modulating the transmural pressure, *11P*, as indicated by the blue arrows. The pressure values associated with the interception points on the abscissa indicate the reopening and the collapse events, which are highly dependent on the surface tension of the liquid film wetting the lumen. Buffer solutions containing bovine liquid extracted surfactant (BLES) at concentrations of 2, 10, and 27 mg ml^-1^ were applied. **Figure 5C** shows a consistent reduction in both the collapse and reopening pressures with decreasing surface tension (i.e., increasing BLES concentration). The finding was the same for the dynamic and the static cases (**Figure S5B** and **S5C**, **Supporting Information**). Furthermore, adding a mixture of BLES and serum protein solutions to the lumen eliminated the reopening pressure reduction (**Figure S5A**, **Supporting Information**). The observation indicates that serum protein may have escaped from the alveolar-capillary interface, causing an elevation in surface tension and impeding the reopening of collapsed airway segments.^[45]^

**Figure 5.**
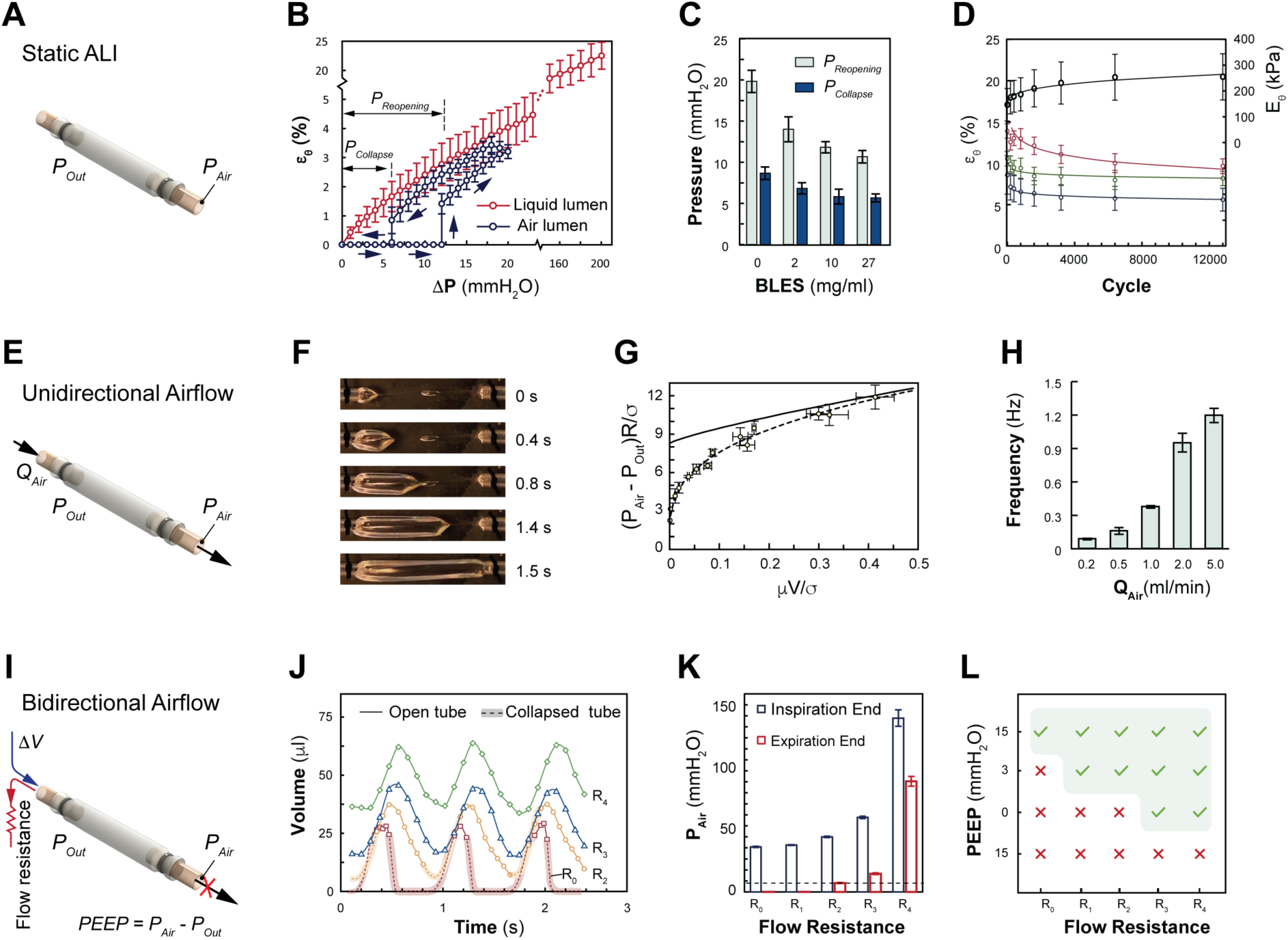
Collapse and reopening of a collagen tube-based airway-on-chip model during static, unidirectional, and bidirectional airflow. (A) Schematic of ALI applied to airway-on-chip model for static airflow. (B) Transmural pressure applied to cannulated collagen tube and corresponding circumferential strain with liquid-filled lumen (**⎯**) and ALI (**⎯**). Arrows (**→**) indicate reopening and collapse events under static ALI. Reopening and collapse pressures determined from abscissa intercepts. (C) Collapse and reopening pressure measured for different surfactant concentrations applied to lumen. (D) Circumferential strain during cyclic stretching performed for pressure amplitudes *P_PEAK_*: 35 mmH_2_O (**—**), 62.5 mmH_2_O (**—**), and 90 mmH_2_O (**—**). Corresponding circumferential Young’s modulus (**—**). (E) Schematic of ALI applied to airway-on-chip model for unidirectional airflow. (F) Time-lapse sequence for reopening of initially collapsed collagen tube subjected to continuous unidirectional airflow. (G) Dimensionless pressure-velocity relationship for collagen tube based on present data (•) and associated fit (--), and Gaver *et al* (-). (H) Collapse and reopening frequency under different airway perfusion flowrates. (I) Schematic illustration of bidirectional ventilation of collagen tube-based aiway-on-chip model with independent control over tidal volume, transmural pressure, and expiratory flow resistance. (J) Tube luminal volume during ventilation with different flow resistances, *R*, at PEEP = 0 Pa, for open (-) and collapsed (--) tubes. (K) Collagen tube luminal air pressures during ventilation at end of inspiration and expiration. (L) Ventilation for wide range of PEEP levels and expiratory flow resistances for collagen tube maintaining open lumen (✓) represents and undergoing repetitive collapse and reopening (✕). Graphs represent mean ± standard deviation, N = 4-5.

Furthermore, long-term cyclic stretch was applied to chip hosted collagen tube segments. During static ALI conditions (i.e., no air flow), the transmural pressure *ΔP* was periodically altered between zero and a maximum value, *Ppeak*, with a frequency of 0.2Hz, as shown in **Figure S3F** (**Supporting Information**). The circumferential strain values and Young’s moduli of the cyclically stretched collagen tube segments were recorded and analyzed (**Figure 5D**) for values of *Ppeak* = 35, 63 and 90 mmH_2_O, and 12,800 cycles. During the initial cycles the circumferential strain decreased significantly and after 6,400 cycles only modestly. The strain reduction between cycles 6,400 and 12,800 was less than 2.2%. The circumferential Young’s moduli increased approximately 50%. These observations suggest the capacity of collagen tubes to be subjected to cyclic stretching at physiologically relevant conditions, while their wall gradually stiffens due to the progressing collagen fibers compaction.

**Figure 5E** shows a schematic representation of continuous unidirectional airflow through the lumen of the collagen tube, with independent control over flowrate and transmural pressure. At *ΔP = 0*, the collapse of the collagen tube was induced by capillary pressure. **Figure 5F** shows a representative sequence of images during reopening an initially collapsed tube by propagating an air finger. The propagation velocity and the pressure were quantified during reopening at a low capillary number, *Ca* = *μV*/*σ* < 0.5, as shown in **Figure 5G**. The experimental data exhibited a favorable fit to the dimensionless relationship described by the equation:

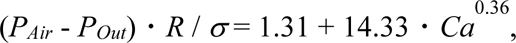

where *R* is the inner radius of the collagen tube after reopening, and *σ* is the surface tension of the liquid film lining the lumen of the collagen tube.

Predictions from a model by Gaver *et al.* 1990^[46]^ are also included in **Figure 5G** for comparison. While our data align with their predictions at high capillary number, a noticeable difference is observed at low capillary numbers (*Ca* < 0.1) that we. We attribute this difference to local changes of the tube diameter close to the upstream suture location. At Δ*P* = 0, shortly after reopening, the soft collagen tube experienced another collapse induced by the capillary pressure. The frequency of this repetitive collapse and reopening phenomenon was found to be influenced by the transmural pressure Δ*P* and the air flowrate (**Figure 5H**).

Subsequently, bidirectional air flow was established within the collagen tube by connecting a miniature ventilator to the air perfusion inlet (Portal I) of the hosting device. The tidal volume, Δ*V* = 30 μL, the positive end-expiratory pressure (PEEP), and expiratory flow resistance, *R_0_* ∼ *R_4_* (for detail see **Table S2**, **Supporting Information**), were independently regulated (**Figure 5I**). PEEP and expiratory flow resistance collectively affected the overall shape and configuration of the collagen tube-based airway model during ventilation. **Figure 5J** shows the variation of the luminal volume of the collagen tube during ventilation at PEEP = 0. Notably, when A small flow resistance (*R_0_*, *R_1_*, or *R_2_*) led to repetitive collapse and reopening. When a higher flow resistance (*R_3_* or *R_4_*) was selected during expiration, tube collapse was avoided, and instead, cyclic stretching was observed. **Figure 5K** illustrates the *in situ* measured luminal air pressures at the end of inspiration and expiration. The collagen tube remained open when both pressure values exceeded the capillary pressure (--). **Figure 5L** summarizes the tube configuration for various PEEP levels and respiratory flow resistances. The findings suggest that increasing PEEP and respiratory flow resistance during the expiration can effectively prevent cyclic collapse and reopening of small airway and alveolar units during the ventilation.

### 2.5. Ventilation Induced Lung Injury

To investigate the conditions related to ventilation induced lung injury in our collapsible collagen tube model, A549 cells were seeded to the lumen, and a confluent layer was obtained after 3 days of culture (**Figure 6A** and **6B**). Subsequently, by varying the tidal volume (Δ*V*) and the flow resistances that are applied in line (*R_0_* to *R_3_*) during bidirectional airflow through the lumen (**Figure 5I**), five distinct ventilation conditions were established as shown in **Figure 6C** and **Table 2**. In condition *i*, *ii*, and *iii*, we successfully replicated the repetitive collapse and reopening phenomena. Capillary numbers were quantified for both the collapse and reopening phases (**Figure 6D**), indicating that a higher flow resistance led to a slower tube collapse during expiration. In the case of overdistension, circumferential strains were recorded when two different tidal volumes were applied during inspiration. Circumferential cyclic stretching with strain levels of 20% and 35% was applied (condition *iv*, *v*, **Figure 6E**). A549 cell-lined collagen tubes were subjected to 50 continuous breathing cycles. **Figures 6F** and **6G** show live-dead staining images and viability. According to our results, repetitive collapse and reopening of our collapsible tube model induces acute cell death, with a viability that decreased for an increasing rate of collapse (corresponding to a larger collapsing capillary number, **Figure 6D**). In contrast, overdistension did not result in significant cell death. No apparent cell detachment was observed in any of the five conditions, suggesting favorable adhesion of the epithelium to the collagen substrate. Furthermore, the IL-8 concentration, cytoplasmatic YAP and permeability of epithelium (using 4kDa dextran) were investigated (see also **Section S7**, **Supporting Information**). Interestingly, both collapse and reopening, as well as overdistension conditions activated the secretion of IL-8 (**Figure 6H**). However, no significant alteration was observed in the immunohistochemical stains for cytoplasmic YAP compared with the control group (**Figure 6I**). Collapse and reopening compromised the integrity of the barrier as indicated by an increased permeability (**Figure 6J**), while overdistension did not result in a noticeable change. These findings suggest that the collapse and reopening phenomena induce acute injury, as opposed to the overdistension during mechanical ventilation. Moreover, in addition to the benefit of expiratory flow resistance in mitigating the occurrence of collapse and reopening phenomena as well as the rate of collapse, our data suggest a reduced magnitude of the injury.

**Figure 6.**
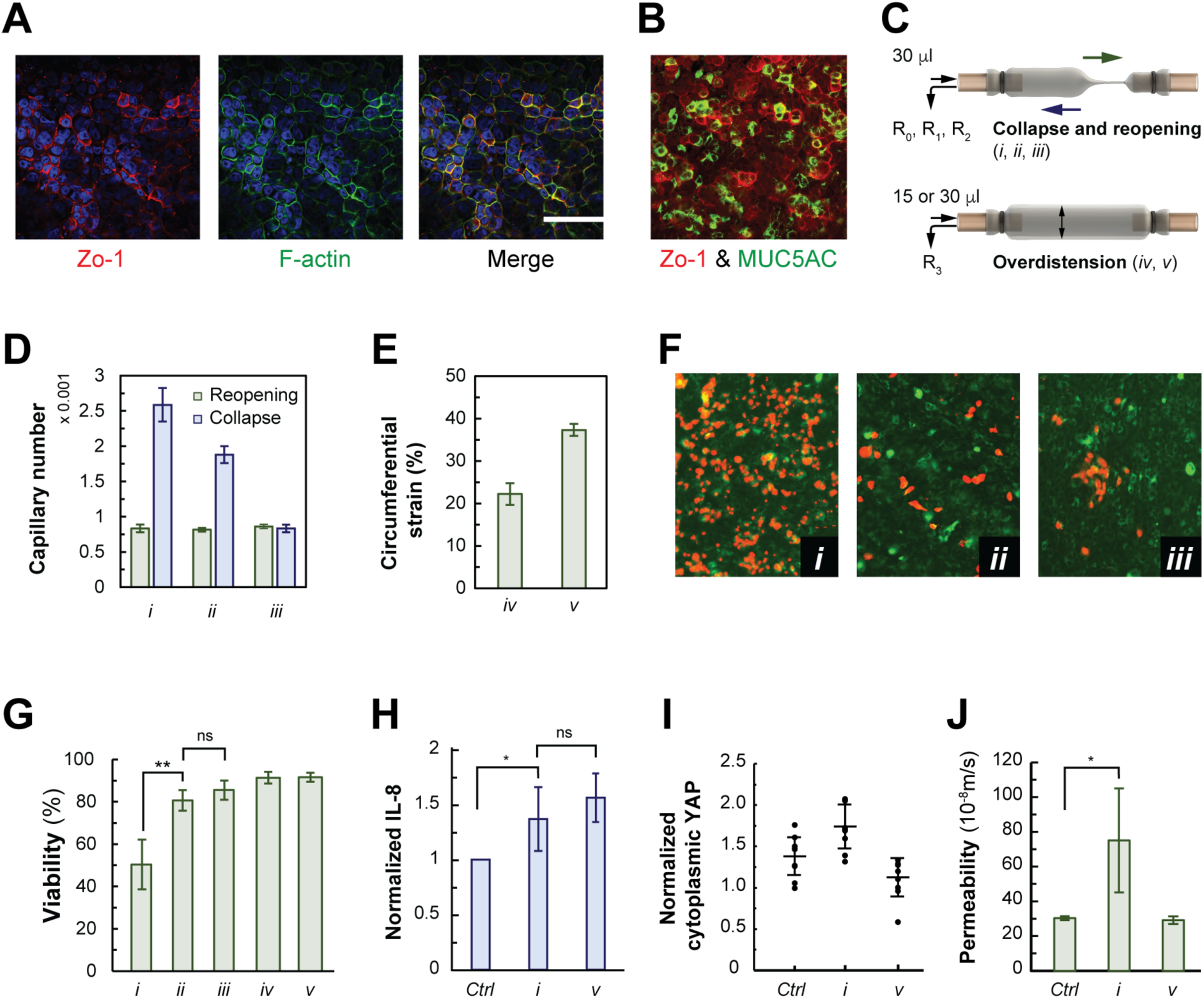
Collapse and stretch induced in collagen tube-based airway-on-chip model at conditions relevant to ventilation induced lung injury. (A, B) Immunostaining images of confluent A549 epithelial cells on lumen of collagen tube. (C) Schematic illustrations of repetitive collapse and reopening (top) and overdistension (bottom) induced in airway-on-chip model. (D) Measured collapse and reopening capillary number for flow resistances *R_0_*, *R_1_*, and *R_2_* applied during expiration. (E) Measured circumferential strain for two tidal volume conditions during ventilation, particularly for flow resistance value *R_3_*. (F) Live-dead staining of A549 monolayer after 50 collapse-and-reopening cycles, for flow resistance values *R_0_*, *R_1_* and *R_2_*. (G) Viability data for collapse and reopening (*i*, *ii*, and *iii*) and overdistension (*iv*, *v*) following 50 cycles. (H) Normalized IL-8 concentration, (I) normalized cytoplasmatic YAP, and (J) A549 epithelium permeability after 50 collapse and reopening cycles (*i*) and overdistension (*v*). Graphs represent mean ± standard deviation, N > 3. ** p<0.01, * p<0.05.

**Table 2.**
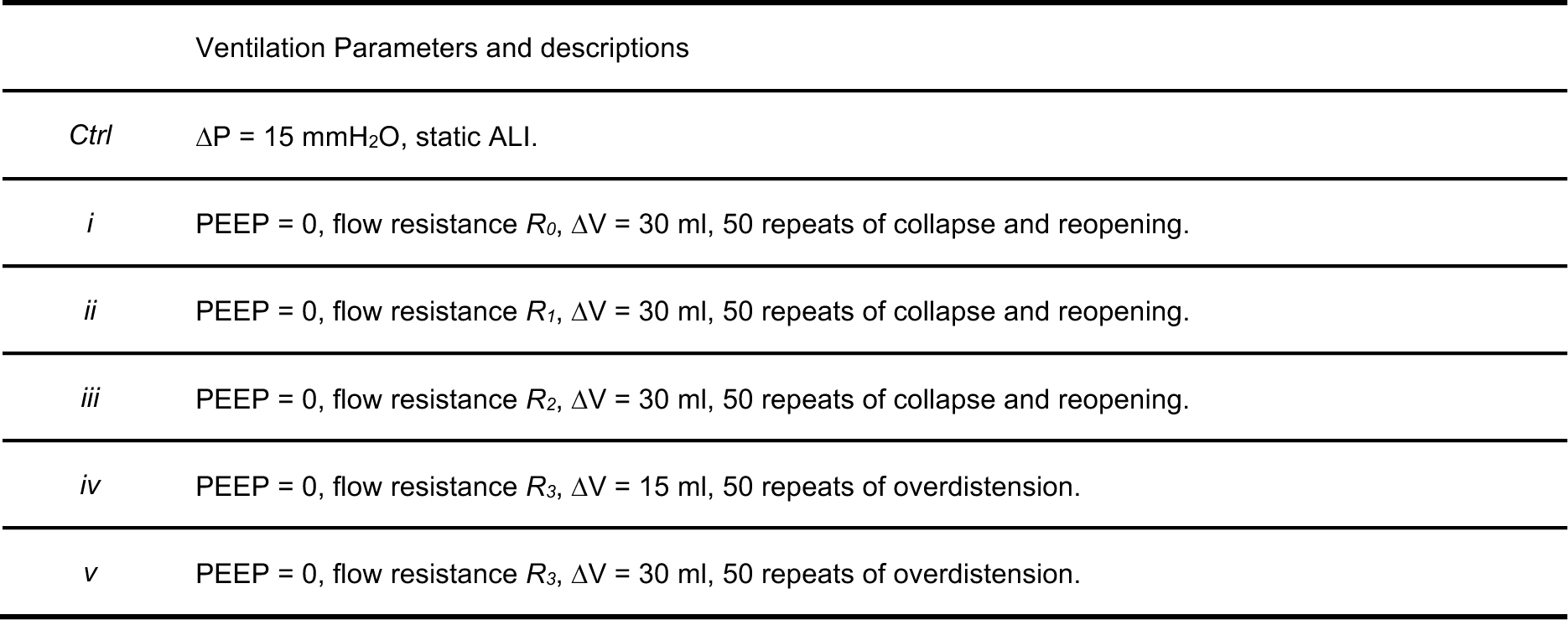
Ventilation conditions considered in collagen tube-based airway-on-chip model.

## 3. Conclusion

In this study, we have presented an airway-on-chip model that is based on an epithelium-lined soft collagen tube. A custom designed tube hosting device facilitates independent perfusion and superfusion, provides *in situ* optical access for inverted imaging at magnifications of up to 20X, promotes the attachemnet of epithelial cells at a high (>90%) yield and culture under both medium and physiologically relevant air flow conditions.

Collagen tube-based airway models exhibit a wide range of tensile properties (e.g., Young’s moduli between 50 kPa and 15 MPa). Leveraging its stretchable and collapsible nature, our model faithfully recapitulates key mechanical features of the airway and alveolar units, including wall shear stress, cyclic stretching, overdistension, and the collapse and reopening phenomena. Specifically, (1) our experiment revealed the distensible collagen tubes to endure up to 12,000 cycles stretching under physiologically relevant conditions. However, we observed an expected decline in the elasticity of the tube wall with an increasing number of cycles. (2) At the ALI, our observations revealed the presence of lung surfactants in the luminal lining fluid to prevent collapse. (3) Additionally, during the mechanical ventilation, elevating either PEEP or air flow resistance (during expiration) had a similar effect.

To evaluate the capacity of our collagen-tube based airway model to recapitulate tissue function and injury conditions, we presented two case studies. In the first case study, a confluent human bronchial epithelium was ALI cultured by careful adjustment of air flow conditions. We found evaporation from luminal lining liquid to be inevitable even for temperature and humidity controlled (37°C, RH > 95%) air flow. Maintaining a consistent osmotic pressures in the culture medium proved critically important for the culture of human bronchial epithelial cells at the collagen tube lumen for up to two weeks under dynamic ALI conditions, validated by high cell viability and enhanced barrier function. Future work may include extending the culture period even further up to 4-5 weeks under bidirectional airflow conditions for iPSC-derived or primary human epithelial cells and coculture with other pulmonary cells.^[11]^ Additionally, extending the range of compatible shear stresses may provide further insights into recapitulating the mechanical microenvironments beyond the presently addressed generations 19 and 20 to the entire respiratory tree, from the trachea through the bronchial passages all the way to the alveolar units.

In the second case study, to our knowledge for the first time, we investigated the key characteristics during the “compliant” collapse and reopening and downstream injuries. Compared to the published findings obtained from “meniscus formation” condition, we obsvered no apparent detatchment of the epithelium and improved cell viability. Our findings also highlight the positive impact of expiratory flow resistance in avoiding or reducing collapse and subsequently reducing lung atelectasis. By adjusting the expiratory flow resistance and the inspiratory tidal volume, our findings suggest collapse and reopening phenomena to lead to acute cell death and loss of epithelial barrier integrity, where overdistension showed no significant effect. Future biomolecular characterization of injuries incurred collagen-tube based airway models lined with patient-specific epithelia may provide a tool to inform patient specific VILI treatment conditions and predict outcomes.

Overall, our soft 3D airway model provides a valuable and adaptable platform for studying lung physiology, particularly ventilation-induced lung injury. The ability to replicate the complex biomechanical microenvironment of the respiratory system enhances our understanding of small airway epithelial function and disease progression, especially for VILI. We consider the presented model to have broad applicability in enhancing human organ-on-a-chip models of different tubular tissues.

## 4. Experimental Section

*Microfluidic hosting device*: the multi-layer thermoplastic hosting device with microchannel networks was fabricated via laser cutting and thermal bonding of acrylic sheets with specific thicknesses. A comprehensive depiction of the tube hosting device design and fabrication process can be found in Section S2, Supporting Information.

*ALI on the collagen tube*: A custom-designed system was developed facilitating controlled air perfusion and culture medium superfusion. Detailed setup components and assembly information is enclosed in Section S3, Supporting Information. Additionally, mechanical simulation of the soft collagen tube under ALI is illustrated in Section S4, Supporting Information.

*Soft tube collapse and reopening*: detailed information of the characterization of the compliant collapse process and parameter settings for the bidirectional airflow within the collagen tube is included in Section S5, Supporting Information.

*Cell seeding, culture, and bioassays*: coating strategy of collagen tube lumen, cell seeding strategy, and cell culture under liquid lumen as well as air lumen are included in Section S6 and S7, Supporting Information.

## Supporting Information

Supporting Information is available in a separate file.

## Acknowledgements

We thank Dr. Nima Vaezzadeh for contributions toward the chip hosting of collagen tubular structures and developing protocols for epithelial cell seeding and long-term culture. We thank Dr. Siwan Park from Dr. Edmond W.K. Young’s laboratory (UofT) for discussions on the air perfusion system, and Dr. Dan Voicu (CRAFT, Toronto) for discussions on device fabrication and feedback on the manuscript, Dr. Keith J. Morton (National Research Council of Canada, NRC, Boucherville, QC) for discussions on device fabrication and Dr. Arthur S. Slutsky (St. Michael’s Hospital, Toronto) on lung surfactants. We thank Dr. Golnaz Karoubi’s laboratory (University Health Network, Toronto) for discusions regarding immunohistochemistry (YAP). APW, AG, CEB, TJM and TV acknowledge support from a CRAFT Research Project Award. AG acknowledges support from NSERC (RGPIN-2017-06781) and the Toronto COVID-19 Action Initiative (TCAI). Device fabrication was carried out at the CRAFT Device and Tissue Foundries that was established with support from the University of Toronto, NRC (Disruptive Technology Solutions for Cell and Gene Therapy Challenge program) and the Canada Foundation for Innovation and the Ontario Research Fund (Ontario-Québec Center for Organ-on-a-Chip Engineering, Center for Advancing Neurotechnological Innovation to Application).

