## Supporting_Text for "Collagen Tubular Airway-on-Chip for Extended Epithelial Culture and Investigation of Ventilation Dynamics"

### S1. Review on Airway on a Chip Platforms

### Table S1. Information of published airway-on-chip (*in vitro*) platforms

| | Target tissue | Platform type | Airway channel | Cells | Airflow | Remarks |
|---|---|---|---|---|---|---|
| 1 | Small airway | Single channel between top and bottom confining cover glass[1,2] | Rectangular (width 10 mm, depth 0.5~1.7 mm) | Rat pulmonary epithelial cell line (CCL-149) | Air and PBS occlusion segmented flow at 0.3 mm/s. | 1. Higher pressure/stress gradient decreased cell viability. 2. Repeated reopening and sub-confluent conditions contributed to cell detachment. 3. Injury from airway collapse and reopening determined from epithelial morphology and focal adhesion. 4. Cells cultured on stiff substrates exhibited more plasma membrane rupture but significantly less detachment and monolayer disrupture. |
| 2 | Large airway | Trans-well insert.[3–5] | Planar (well plate insert diameter 6 mm) | Primary human bronchial epithelial cells | N/A | 1. Ciliated surface area ratio reached 75% after 3-week ALI culture. 2. Swirls of mucus flow observed within well plates insert, 1.3 mm and 0.6 mm in size. 3. Cilia beating direction mostly consistent with mucus swirl. 4. Mucus removal and dehydration found to induce random cilia beating. |
| 3 | General airway | Modified well-plate insert model.[6] | Planar (well plate insert diameter 12 mm) | Primary human tracheal epithelial cells, human lung fibroblast cell lines, and vascular endothelial cell lines. | N/A | 1. Sophisticated trans-well model demonstrates asthmatic airway inflammation and allergen-induced asthma exacerbation. |
| 4 | Small airway | 2 PDMS channels separated by polyester membrane (0.4 µm pores)[7,8] | Rectangular (width 0.3 mm, depth 0.1 mm) | Primary human small airway epithelial cells (SAECs) | Air flow and PBS liquid plug occlusion, segmented flow at 1.5~5 mm/s. | 1. Propagation and rupture of liquid plugs mimicking surfactant deficient reopening of occluded airway led to significant epithelial cell injury. 2. Pressure wave from plug rupture produced crackling sound that correlated with cellular injury and had similarities with clinical observations. |

| | | | | | | |
|---|---|---|---|---|---|---|
| 5 | Distal airway | 2 PDMS channels separated by polyester membrane (0.4 µm pores) [12] | Rectangular (width 1 mm, depth 0.5 mm) | Primary human small airway epithelial cells | 1.7 mm/s PBS exudated form collagen gel serves as mucus plug. | 1. Propagation of mucus plugs in airway was recapitulated. |
| 6 | Small airway, general airway (bronchiole) | 2 PDMS channels separated by polyester membrane (0.4 µm pores) [9–11] | Rectangular (width 1 mm, depth 1 mm) | Primary human airway epithelial cells and human lung microvascular endothelial cells seeded onto opposing sides of membrane. | N/A | 1. Small airway on chip model containing a differentiated, mucociliary bronchiolar epithelium and microvascular endothelium.<br>2. Exposure of epithelium to IL-13 induced goblet cell hyperplasia, cytokine hypersecretion and decreased ciliary function.<br>3. Model with patient specific cells recapitulated disease characteristics.<br>4. Supply of cigarette smoke allowed comparison of cellular responses with/without smoke exposure.<br>2. Viral infection, strain-dependent virulence, cytokine production and recruitment of circulation immune cells demonstrated. |
| 7 | Large airway | 2 thermoplastic channels separated by suspended hydrogel (Matrigel & collagen, 0.65 mm thick) [12] | Rectangular (width 2 mm, depth 1.9 mm) | Human airway epithelial cells (Calu-3) and human bronchial smooth muscle cells seeded onto opposing sides of hydrogel. | ~ 10 mm/s | 1. Airway epithelial cells displayed cobblestone morphology with tight junctions, positive for MUC5AC staining under ALI.<br>2. Long-term viability during SMC-SMCs coculture.<br>3. Enable complete sample extraction and off-chip analysis.<br>5. ALI culture is limited to durations shorter than 3 days. |
| 8 | General airway | 3 PDMS channels separated by PTFE and PET membranes [13] | Rectangular (width 1 mm, depth 0.28 mm) | Human trachea-bronchial epithelial cells, primary human lung fibroblast, and human lung microvascular endothelial cells | N/A | 1. Mucuciliary differentiation and barrier function (permeability of dextran 20) was demonstrated with three cell type coculture. |
| 9 | Small airway | Casted collagen | Circular | Pulmonary fibroblast cell | N/A | 1. Recapitulate human bronchiole by coculturing co-located |

| | | | | |
|---|---|---|---|---|
| (bronchiole) | (6 mg/ml) channel with extractable PDMS rod.[14] | (diameter 0.55 mm) | line, Primary human bronchial epithelial cells, and human lung microvascular endothelial cell line | airway epithelium, pulmonary fibroblast, and endothelial cells.<br>2. Communication of volatile compounds between microbial populations and host model demonstrated. |

### S2. Tube Hosting Device Design and Fabrication

**Figure S1A** shows a rendered image of assembled tube hosting device with manually cannulated collagen tube. **Figure S1B** illustrates the perfusion and superfusion strategy within the tube hosting device, including perfusion inlet (I), outlet (II or III), superfusion inlet (IV), outlet (V or VI). **Figure S1C** illustrates a 20× magnification optical access of bottom lumen surface of a sutured collagen tube. **Figure S1D** shows the cell seeding reservoir design between portal II and III with channel surface coated by anti-adherence rinsing solution (Stemcell Technologies, Vancouver, BC, Canada). A reservoir with a volume of 75 µl was included underneath portals II and III to allow for cell seeding. **Figure S1E** schematically illustrates the fabrication process of the tube hosting device via laser cutting and thermal bonding of plastic sheets (Clear cast acrylic sheets with varying thicknesses: Layer 1 - 0.2 mm, Layer 2 - 0.8 mm, Layer 3, 4, 7, 8 - 1.6 mm, and Layer 5, 6 - 3.2 mm. From McMaster-Carr in IL, USA.). Fluidic channels in the different feature layers were designed in AutoCAD (2022, AUTODESK, San Rafael, CA, US). A laser cutter (model VLS3.60, Universal Laser Systems, Scottsdale, AZ, US) with a 10.6 µm wavelength carbon dioxide laser source was used to cut layers (1-8) of the tube hosting device. A circular hole (diameter 1.3 mm) was cut in the cannula adapters VII to accommodate the plastic dispending needles.  A groove was laser machined into lid IV (indicated in dark gray color) to accommodate a custom molded PDMS O-ring.

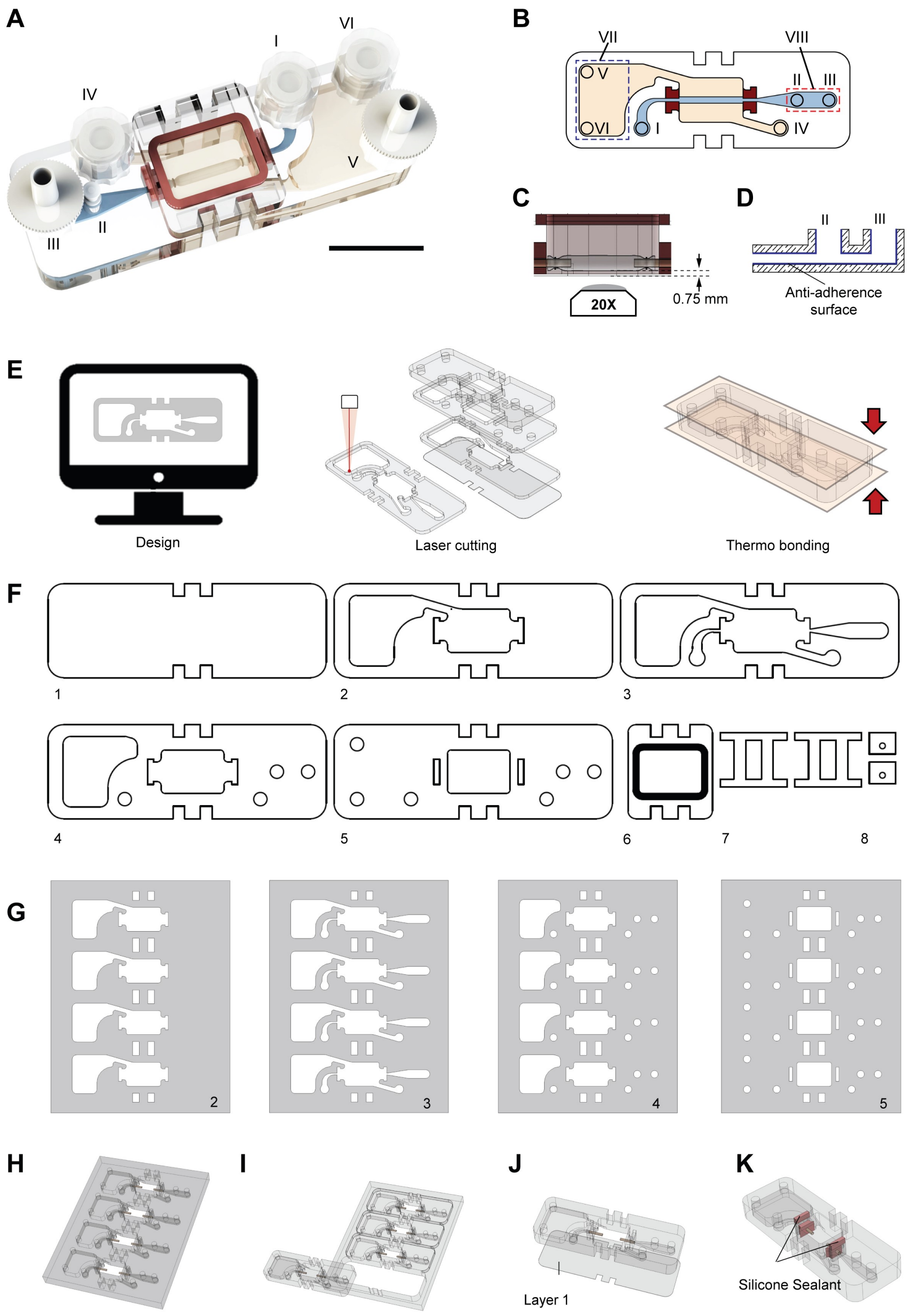


**Figure S1. Schematic of collagen tube hosting device fabrication.** (A) Rendered image of assembled tube hosting device. Scale bar: 2 cm. (B) Schematic illustration of perfusion and superfusion strategies in tube hosting device. (C) Illustration of optical access to 20× magnification inverted microscope objectives to the bottom lumen surface of cannulated collagen tubes. (D) Design of cell seeding reservoir (75 μL). (E) Illustration of fabrication sequence including design, laser cutting, and thermal bonding. (F) Flow channel design of tube hosting device with (I) bottom cover layer, (II) - (IV) fluidic channel layers, (V) top layer, (VI) lid with O-ring, (VII) clamps, and (VIII) cannula blocks accommodating dispensing needles. (G) - (K) illustrate simultaneous manufacture (i.e., cutting and bonding) of four devices.

To increase throughput of tube hosting device manufacture, four devices were fit on a 6” wafer footprint. Thermal bonding was applied to all devices simultaneously. Individual tube hosting devices were dissected, and a bottom film (layer 1) made from an acrylic adhesive (**Figure S1I** and **S1J**) was attached. Silicone sealant (Momentive$^{TM}$ New York, USA) was used to seal the gap between horizontal bonded thermal plastic sheets, vertical cannula adapters, and dispending needles. After fabrication, each tube hosting device was submerged in 70% ethanol for 20 min and air dried under UV exposure (~30 mW cm$^{-2}$, Thorlabs Saint-Laurent, QC, Canada) for 20 min in a biosafety cabinet for sterilization.

**S3. Tube Hosting Strategies**

To recapitulate the ALI culture of airway epithelial cells, we provided controlled air perfusion and culture medium superfusion. **Figure S2A** illustrates the transition of the collagen tube-based airway on chip device from a culture medium filled lumen to air-liquid interface culture. In the latter case, air was introduced from portal I, while the syringe filter at portal III was substituted by a G21 syringe needle.

**Figures S2C** and **S2D** show the schematic of this 32-tube culture system. A pressure regulator was connected to a compressed air source (100 PSI). An inline air filter (Nylon plastic, McMaster-Carr, Los Angeles, CA, USA) was employed to retrain particles with sizes exceeding 0.1 μm. Additionally, another inline air filter (0.22 μm pore size, Millipore Sigma, Oakville, ON, Canada) was added downstream of the air pressure regulator. To achieve the desired humidity and temperature levels resembling physiological conditions, the air flow was passed into a heated reservoir filled with deionized water through a porous air-stone. A T-connection, equipped with a valve and a filter, facilitated the addition of de-ionized water. A temperature sensor was connected to monitor real-time the air temperature within the reservoir. Additionally, a second reservoir was connected downstream to collect condensate and prevent water from traveling to the downstream flow resistance channels. A "1-4" manifold (McMaster-Carr, Los Angeles, CA, USA) was used to simultaneously provide consistent airflow to several parallel collagen tube-based airway on chip devices. Each of the four manifold outlets was connected to an 8-device platform (fabricated from thermoplastic) through a 1.5 m long tygon tubing via a valve (McMaster-Carr, Los Angeles, CA, USA) that enables independent transfer of each platform between the tissue culture incubator and a nearby biosafety cabinet for superfusion medium exchange. The 8-device platform contained a series of flow resistors, liquid traps, as well as eight collagen tube-based airway on chip devices (**Figure S2D**). Air was perfused into the individual airway on chip device from Portal I to III, leaving through a downstream syringe needle. To realize physiological air velocity (2.5 cm $s^{-1}$), we select polyether ether ketone (PEEK) tubing (inner diameter 63.5 μm) for the individual air flow resistors. The downstream syringe needle (21G, 25.4 mm long) was connected to provide a small pressure offset that maintained the lumen of the collagen tube open during dynamic ALI culture at the given air velocity. **Figure S2B** shows the simulated air pressure distribution along the airway on chip model while being perfused with an air velocity 2.5 cm/s, when needles with different gauge values were considered. We define the transmural pressure as $\Delta P = P_{IN} - P_{CAP} - P_{OUT}$, where $P_{IN}$ is air pressure in tube, $P_{CAP}$ is the capillary pressure associated with the liquid film on the lumen, $P_{OUT}$ is the pressure in the abluminal culture medium. To change culture medium within the abluminal space, each platform was carefully transferred to biosafety cabinet from incubator while air flow is maintained. Fresh medium was added to Portal IV, and medium was withdrawn from Portal V.

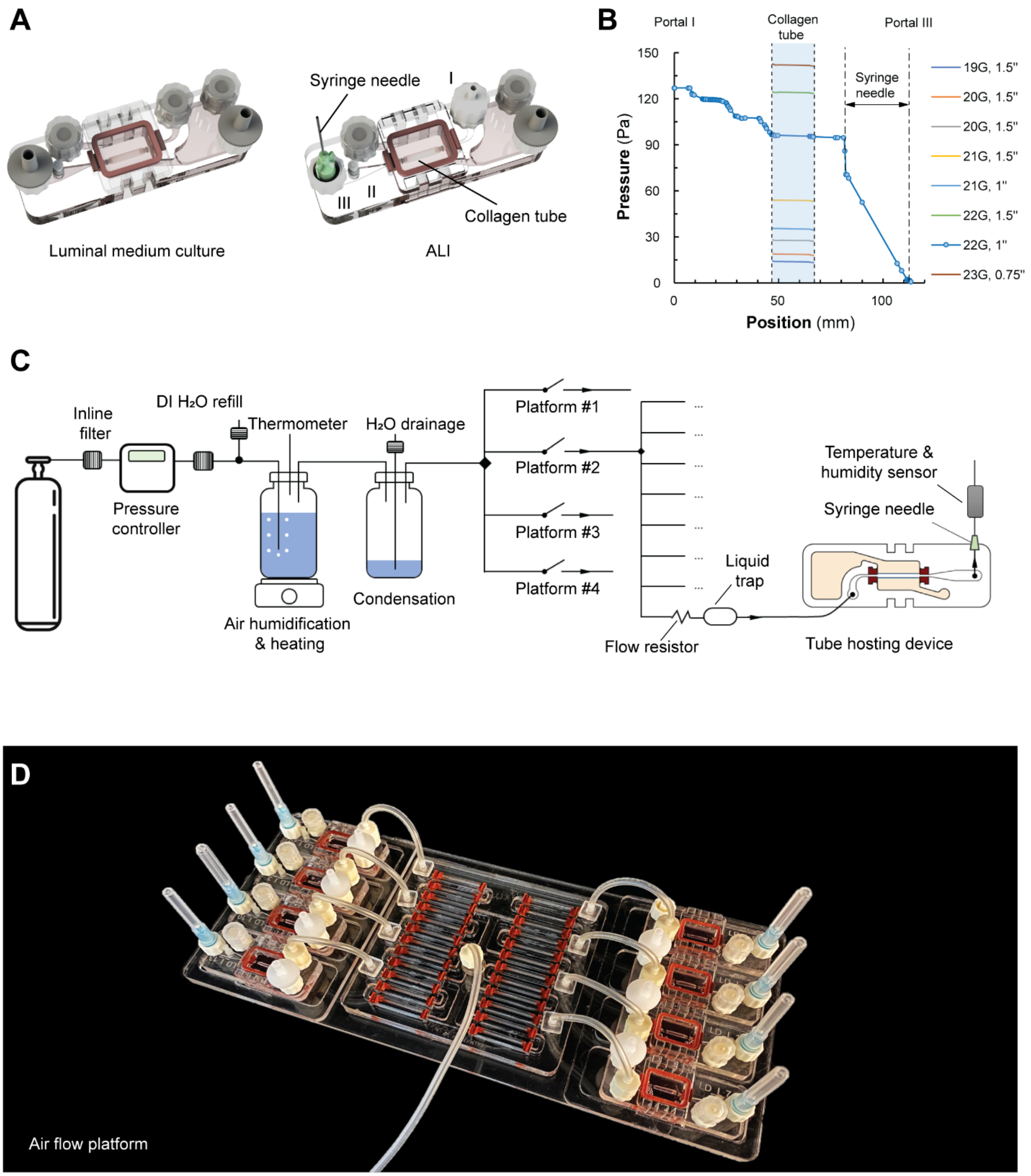


**Figure S2. Strategies for dynamic ALI culture in collagen tube-based airway model.** (A) Schematic of tube hosting device before and during ALI culture. (B) Pressure-drop within tube hosting device along air perfusion streamline. Dark area represents local pressure within collagen tube when applying syringe needle downstream for increased flow resistance. (C) Schematic of airflow system containing air source, pressure controller, air humidification and heating, water condensation and four platforms. (D) Photograph of each individual platform, which can accommodate 8 tube hosting devices in parallel.

**S4. Effects of Buoyancy, Cyclic Stretch, and Wall Shear Stress on Collagen Tube Under ALI Conditions**

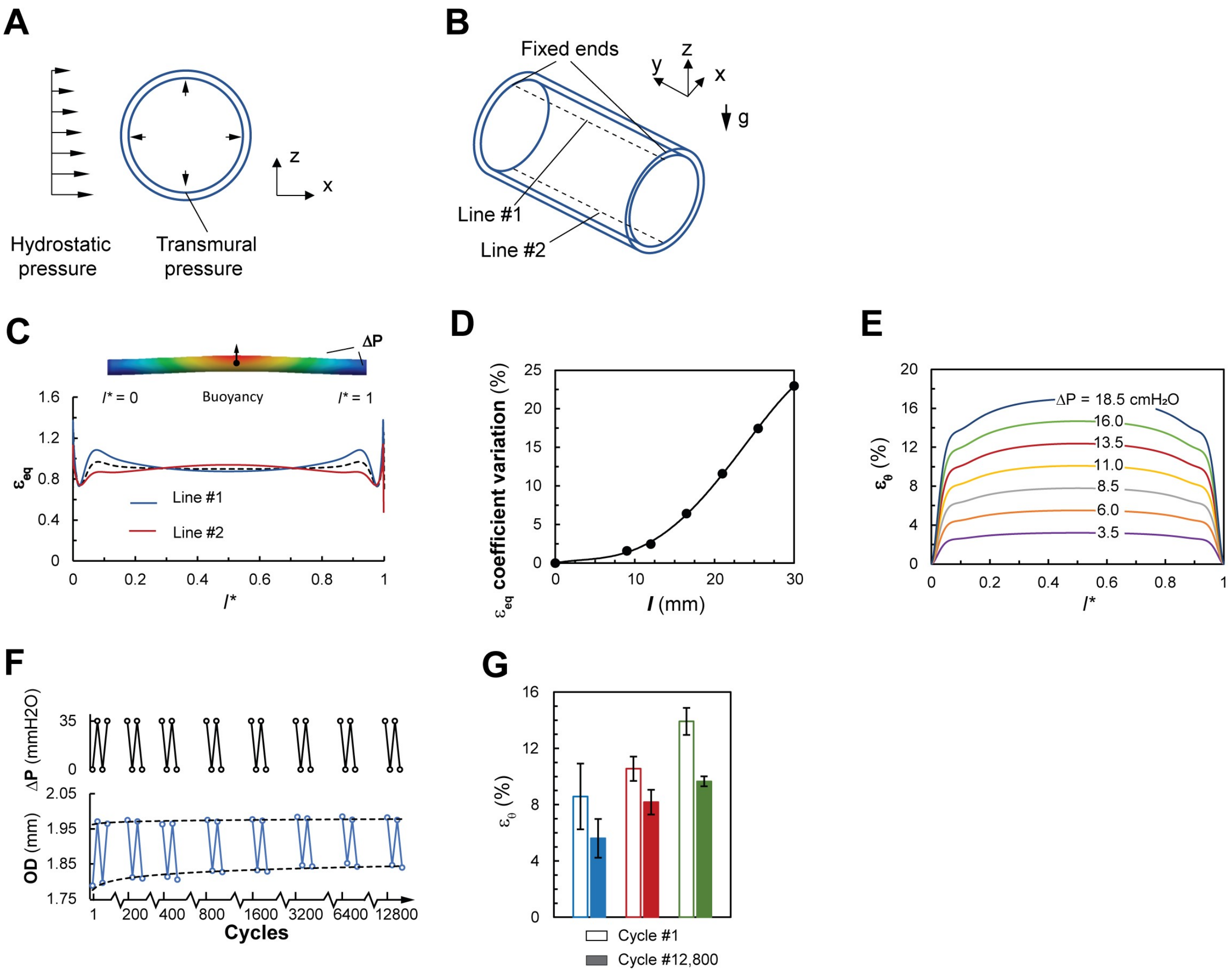


**Figure S3. Cyclic stretching of collagen tube segment under ALI conditions.** (A) and (B) Simulation setting of hydrostatic pressure, transmural pressure, two fixed ends, and two targeting lines for strain distribution analysis. (C) Simulation of buoyancy effect on strain of hosted collagen tube under ALI interface (ΔP = 9.2 $mmH_2O$) along Lines #1 and #2 as indicated in (B). (D) Variation of equivalent strain distribution between top and bottom lumen at ΔP = 3.5 $mmH_2O$ for tube lengths between 3 mm and 30 mm. (E) Simulated circumferential strain distribution along the tube length with varied transmural pressure from 3.5 to 18.5 $cmH_2O$. (F) Example of diameter change for collagen tube during 12,800 cyclic stretching under coded transmural pressure, ΔP = 0 ~ 35 $mmH_2O$, at frequency 0.2 Hz. (G) Cyclic circumferential stretch applied to collagen tubes via peak pressure values, $P_{PEAK}$: 35 $mmH_2O$ (blue), 62.5 $mmH_2O$ (red), and 90 $mmH_2O$ (green).

To assess the buoyancy effects on the cross-sectional shape and the strain distribution for a collagen tube segment that is air-filled and surrounded by culture medium, we conducted a finite element simulation under static conditions using a commercial solver (Academic Research Mechanical, ANSYS, Canonsburg, PA, USA). The three-dimensional geometry of the undeformed cylindrical tube segment was generated using Solidworks and imported into the ANSYS Mechanics environment. The diameter was 1.75 mm, the thickness was 22 μm, and the length varied between 3mm and 30 mm. The material properties for the collagen tube segments were selected as follows: density 998.2 kg/m$^3$, orthotropic elasticity $1.876 \times 10^5$ Pa in the circumferential direction and $2.389 \times 10^6$ Pa in the axial direction, and a Poisson ratio of 0.25[15]. As shown in **Figure S3A**, the two ends of the collagen tube segment were defined as fixed supported, the transmural pressure was applied as a normal force acting on the lumen surface of the tube segment and increased in increments from 3.5 to 18.5 mm$H_2O$, while the buoyancy force acted on the tubular structure.

After convergence, the distribution of the numerically predicted strain was evaluated along the two lines shown in **Figure S3B** (indicated as Line #1 and Line #2, marking the top and bottom on the lumen in the vertical direction, respectively), to assess its uniformity and the effect of buoyancy. In **Figure S3C**, the obtained strain values, $\varepsilon_{eq}$, are plotted for a 12 mm-long collagen tube segment that was subjected to a transmural pressure of $\Delta P$ = 3.5 cm$H_2O$. As a control, the black dashed line represents the distribution for a culture medium filled lumen, while the red and blue lines represent the strain profiles on the top and bottom lines on the luminal surface of the air perfused lumen. The numerical results suggest the strain to vary less than 2.5% within the 3/5 center part of the collagen tube length (i.e., away from the fixation locations), at transmural pressures $\Delta P < 5$ cm$H_2O$. **Figure S3D** shows the coefficient of variation when different tube lengths were considered. To ensure a relatively uniform strain distribution, the length of tube segments considered in our experiments was selected to be between 10 mm and 12 mm. **Figure S3E** shows the numerically predicted circumferential strain values for a wide range of transmural pressures at the ALI, and they agree with the measured data shown in **Figure 5B**.

The airways of the respiratory system are typically subjected to 4% cyclic strain at 0.2 Hz during quiet tidal breathing and 25% strain at 0.43 Hz during sighs or heavy work. We used a linear translation stage to impose corresponding transmural pressures under ALI conditions and reproduce the long-term cyclic stretching phenomenon. Under ALI conditions, the transmural pressure $\Delta P$ was periodically altered between 0 and the peak value, $P_{PEAK}$, at 0.2 Hz, as shown in **Figure S3F**. The resulting cyclic stretch was recorded. In **Figure 5D** and **Figure S3G**, $P_{PEAK}$ = 35, 62.5, and 90 mm$H_2O$ were applied, for up to 12,800 cycles of cyclic stretching. Both circumferential strain and Young's modulus were recorded over time.

Subsequently, the wall shear stress was numerically predicted within the deformed tube geometry. **Figure S4** compares the wall shear stresses on the luminal surface. For a cross-sectionally averaged air velocity of 2.5 cm/s, a uniform wall shear stress of 2.5-2.6 mPa was found along the circumference of the collagen tube. For the same average air velocity applied to a square microchannel, the wall shear stress assumes a maximum value of 2.7 mPa at the center of each wall surface, and a minimum of 0.4 mPa at four corners.

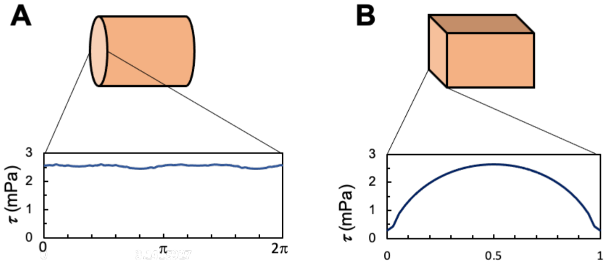


**Figure S4.** Variation of wall shear stress in circumferential direction, for (cross-sectionally) averaged air velocity of 2.5 cm/s, compared between microfluidic channels of (A) circular and (B) square cross section.

### S5. Compliant Collapse and Reopening of Soft Collagen Tube

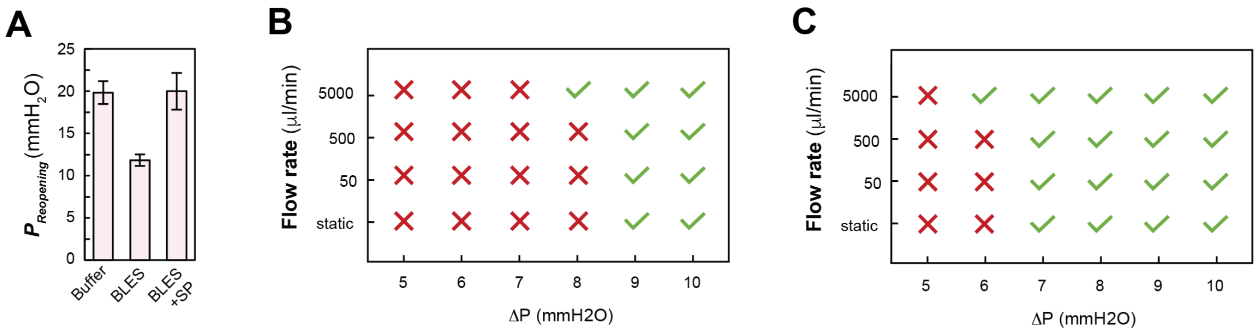


**Figure S5. Collapse and reopening phenomenon with variable air perfusion flowrate.** (A) Reopening pressure of collagen tube with luminal surface lining buffer, 10 mg ml$^{-1}$ BLES, and 10 mg ml$^{-1}$ BLES with serum protein solutions. Tube configuration under varied air flowrates and transmural pressure when luminal surface lined with buffer only (B) and 10 mg ml$^{-1}$ BLES (C).

For the static airflow ALI condition, transmural pressure was adjusted continuously to investigate the tube configuration and assess effect of the surfactant BLES (**Figure 5C** and **Figure S5A**). For the unidirectional airflow ALI condition, air flowrate was precisely regulated by a syringe pump, set at 50, 500, and 5000 μl min$^{-1}$. The tube configuration was evaluated under different transmural pressures and flowrates, in the absence and presence of 10 mg ml$^{-1}$, as depicted in **Figure S5B** and **S5C**, respectively.

To provide bidirectional airflow within the hosted collagen tube, a miniature ventilator (Mini-Vent, type 845, Harvard Apparatus, Holliston, MA, USA) was connected to the perfusion portal (III), while the perfusion outlet was closed. The PEEP (identical to the transmural pressure), the tidal volume, and the expiration flow resistance were controlled by the superfusion pressure (**Figure 5I**). **Table S2** shows the detailed information about the flow resistance applied during the bidirectional airflow.

**Table S2.** Information on flow resistance tube used during the expiration phase of ventilation.

| | Flow resistance tube type, inner diameter, length | Flow resistance (Pa·s/ml) |
|---|---|---|
| $R_0$ | Tygon tube, 1.6 mm, 100 mm | 19.0 |
| $R_1$ | G20 needle, 0.603 mm, 38.10 mm | $2.15 \times 10^2$ |
| $R_2$ | G23 needle, 0.337 mm, 38.10 mm | $2.20 \times 10^3$ |
| $R_3$ | G25 needle, 0.260 mm, 19.05mm | $3.10 \times 10^3$ |
| $R_4$ | G30 needle, 0.159 mm, 12.7 mm | $1.48 \times 10^4$ |

### S6. Development of Epithelium from HBE Cells Seeded to Lumen of Collagen Tube.

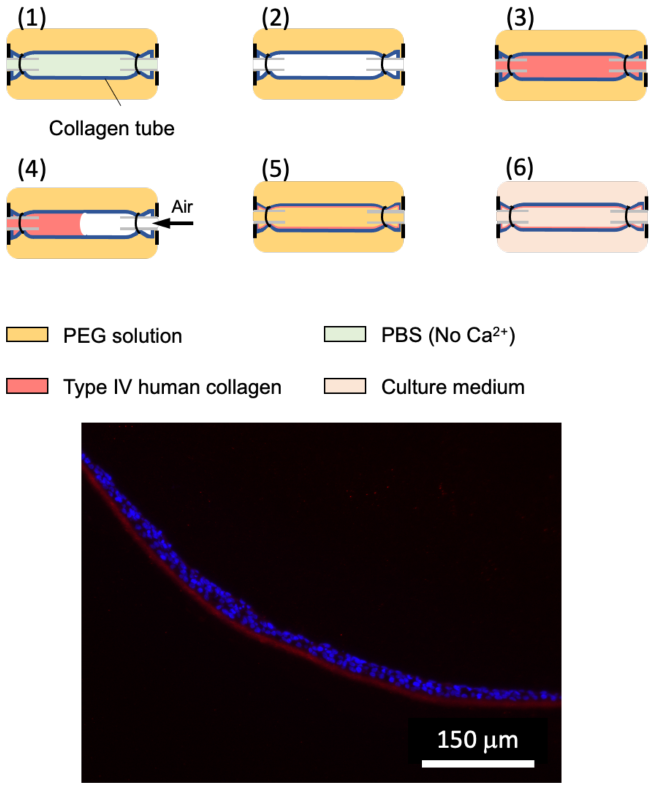


**Figure S6. Coating of Collagen Tube Lumen with Type IV Collagen**

To promote cell attachment during ALI, human type IV collagen (derived from human placenta, Millipore Sigma, Oakville, ON, Canada) was coated on the lumen of the collagen (type I collagen derived from bovine) tube segments. Acidic human type IV collagen solution with a concentration of 0.6 mg/ml was prepared in advance. As shown in **Figure S6**, this coating strategy includes:

(1) Fill the abluminal space with PEG solution ($M_W$ =35,000, 10%, pH 8).

(2) Fill lumen initially with PBS. Then replace by air and maintain for 20 min.

(3) Perfuse 70 µl type IV human collagen solution and maintain for 2 min.

(4) Continuously perfuse air into collagen tube at 0.5 ml/ml for 20 min.

(5) Replenish collagen tube lumen with same PEG solution and maintain for 1 h.

(6) Rinse collagen tube three times with PBS (without $Ca^{2+}$), and twice with culture medium.

(7) Maintain collagen tube open during this process with a positive transmural pressure.

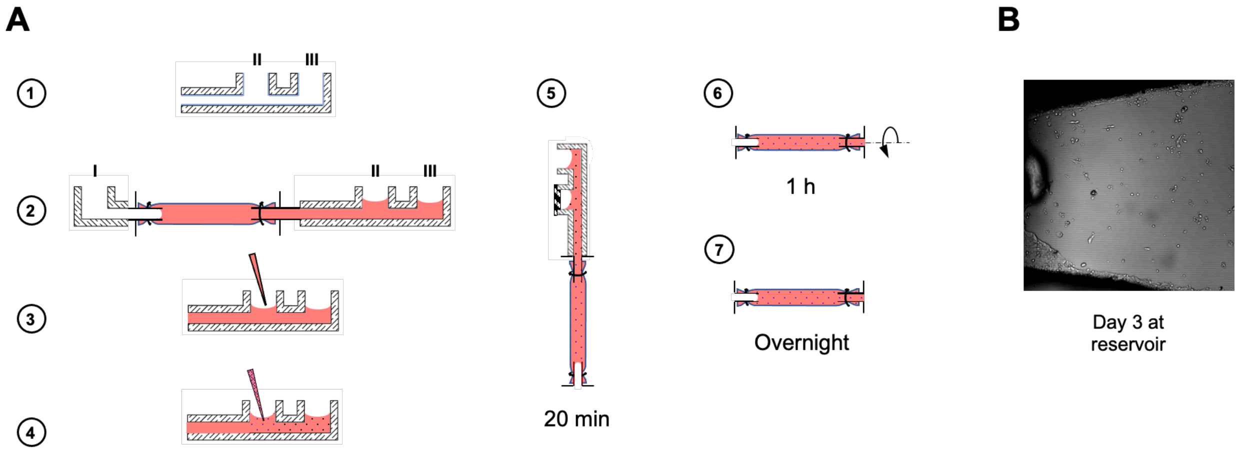


**Figure S7. Cell seeding strategy.** (A) Schematic of cell seeding within the superfusion channels. (B) Image at the cell seeding reservoir at day 3 after cell seeding step.

Cells were seeded into tube hosting device from portal II and III as below:

(1) Before suturing collagen tube, fill the lumen surface of small reservoir beneath portal IV and V with anti-adherence rinsing solution (Stemcell Technology, BC, Canada) for 25 min, then rinse with PBS 3 times.

(2) Fill the reservoir VII with cell culture medium, perfuse air from portal I and stop the air liquid interface at the tip of the plastic needle.

(3) Remove 50 μl culture medium from portal II.

(4) Add 50 ml cell suspension (concentration: 4 or 10 million cells per ml) into portal II. Then seal portal II by straight plug (barbed tube, #5152K121, McMaster-Carr), connect portal III with syringe filter.

(5) Keep device vertical for 20 min, the air liquid interface introduced from step 2 prevents cells settling further beyond collagen tube.

(6) Keep tube hosting device onto a rotating stage,[16] rotate along collagen tube axis at 0.2 rpm for 1.5 h.

(7) Maintain the tubing hosting device in the incubator overnight, then remove the air from portal I.

As illustrated by **Figure S6B**, very few cells were left within the reservoir VII, demonstrating high cell seeding yield.

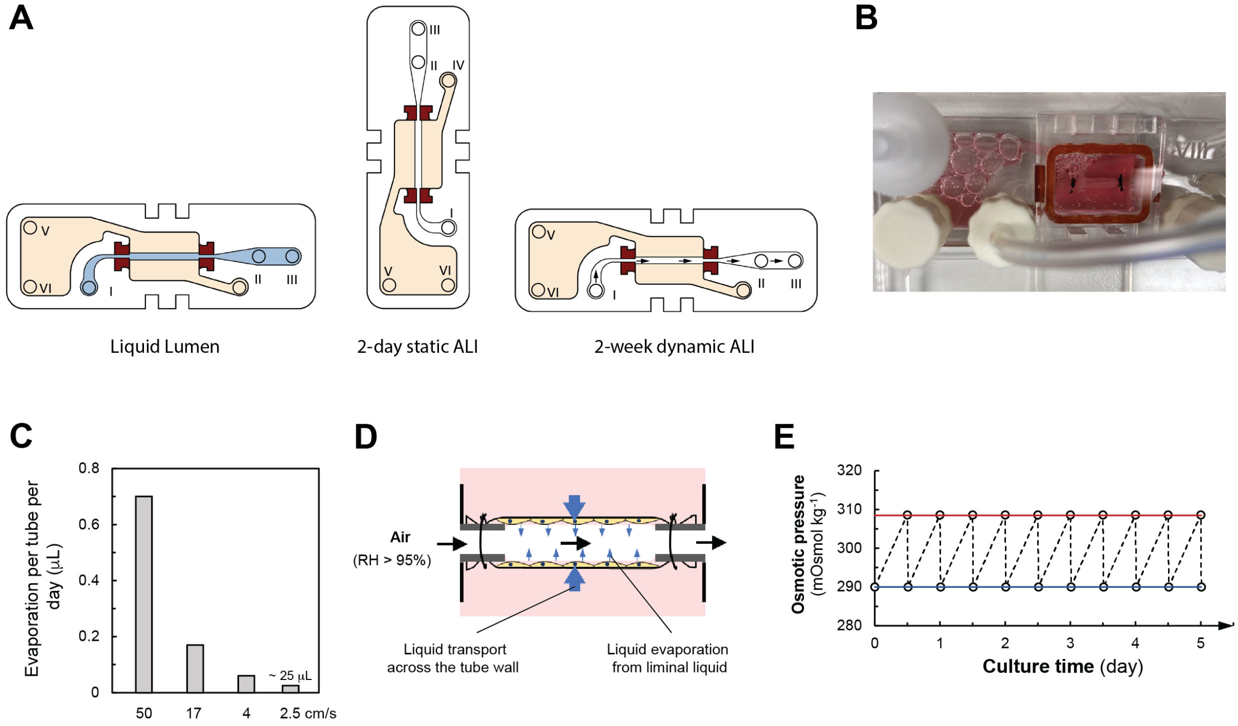


**Figure S8. Cell culture strategies.** (A) Schematic of transition between culture medium filled lumen to dynamic ALI culture. (B) Photograph showing evaporation within superfusion space during air perfusion. (C) Measured volume of evaporated water per device and day, for three different air flow velocities applied at the lumen. (D) Schematic illustration of water evaporation from luminal surface and transport across tube wall. (E) Estimated osmotic pressure in medium on superfusion side during ALI culture (influence of cells neglected) using described approach.

To achieve long term dynamic ALI culture of HBE cells, we have developed a cell culture strategy as illustrated in **Figure S8**.

(1) For a liquid-filled lumen, both perfusion and superfusion channels are filled by EMEM supplemented with FBS (10%) and penicillin/streptomycin (1%). Every other day, 0.5 ml fresh medium was added from portal I, and the same volume of old medium was removed from portal III. Similarly, 2.5 ml medium was exchanged in the superfusion from portal IV to V.

(2) Once confluent epithelium was achieved on the luminal surface, EMEM medium in the superfusion channel was changed by PneumaCult$^{TM}$-ALI medium (Stemcell technology, BC, Canada), supplemented with 1X PneumaCult supplement, 1% v/v maintenance supplement, 0.2% v/v heparin solution, 5 mM dexamethasone solution). EMEM medium in the perfusion channel was removed. While portal IV was closed, tube hosting device was placed vertically to maintain $\Delta P \approx 15$ mm$H_2O$ for 48 h in the incubator.

(3) To establish dynamic ALI, portal III was connected to a G21 syringe needle, meanwhile portal I was connected to the air flow platform. Every other day, differentiation medium was exchanged in the superfusion channel from portal IV to portal V. Note that the transmural pressure across the tube is small, ~ 92 Pa, careful manipulation is necessary to transfer platform from the incubator to the biosafety cabinet, and medium exchange in the superfusion to maintain positive transmural pressure, preventing unwanted tube collapse and reopening. Also, during air flow, evaporation occurred within the luminal liquid (**Figure S8B ~S8D**), due to the relative humidity in the conditioned air flow not exactly matching 100%. The medium in the superfusion was distributed across the tube wall by capillary force. We found approximately 25 μL water to evaporate from the medium every

day when epithelium was exposed to airflow at average velocity at 2.5 cm/s per day, causing an unwanted change in the osmotic pressure of the culture medium. Warm distilled water was added twice a day from portal VI into reservoir VI, to maintain initial osmatic pressure in the superfusion. Figure S7E shows the calculated osmotic pressure alternation during the culture.

### Section S7. Epithelial Tissue Characterization

#### FITC-Dextran Flux Assay

FITC dextran (Mw 4,000) was selected to analyze the permeation of macromolecules across the collagen tube wall.[17] For **Figure 4M** and **6J**, FITC dextran solution at constant concentration $C_0$ = 0.2 mM was added into the perfusion channel. The pressure at the luminal and abluminal side were kept at $\boldsymbol{P_{ATM}}$. Every 5 min, a 0.5 ml sample was collected from the ablumen (total volume, $V_{AL}$ = 2.6 ml) and evaluated using a fluorescence spectrometer (excitation 483 nm, emission scan 518 ± 10 nm, Cary Eclipse, Agilent Technologies), and returned. Over time, $t$, the transport of dextran across the wall of the collagen tube segment with the lumen surface area, $A$, and tube wall thickness, $l$, can be described as below:

$$V_{AL}\frac{dC_{AL}(t)}{dt} = \frac{AK_dD}{l}[C_L(t) - C_{AL}(t)], \qquad \text{(S1)}$$

$$(C_0 - C_L(t)\,) \cdot V_L = C_{AL}(t) \cdot V_{AL}, \qquad \text{(S2)}$$

where $C_L$ and $C_{AL}$ are the dextran concentrations in the tube lumen and ablumen spaces, respectively. $V_L$ and $V_{AL}$ are the volumes of tube lumen and ablumen space, respectively. $K_d$ and $D$ are the partition coefficient and the diffusion coefficient for molecular transport across the tube wall, respectively.

The permeability coefficient for the collagen tube wall is defined as below:[18]

$$P = \frac{K_d D}{l}. \tag{S3}$$

Applying Equations (S2) and (S3), and the collagen tube dimensions, Equation (S1) is calculated.

$$C_0 - 11.13 \cdot C(t) = h_1 \cdot exp(-2.87 \times 10^2 \cdot P \cdot t) + h_2, \tag{S3}$$

Using Equation (S3), the permeability coefficient (**Figure 4M**) was calculated from the time-dependent concentration data (**Figure S9B**).

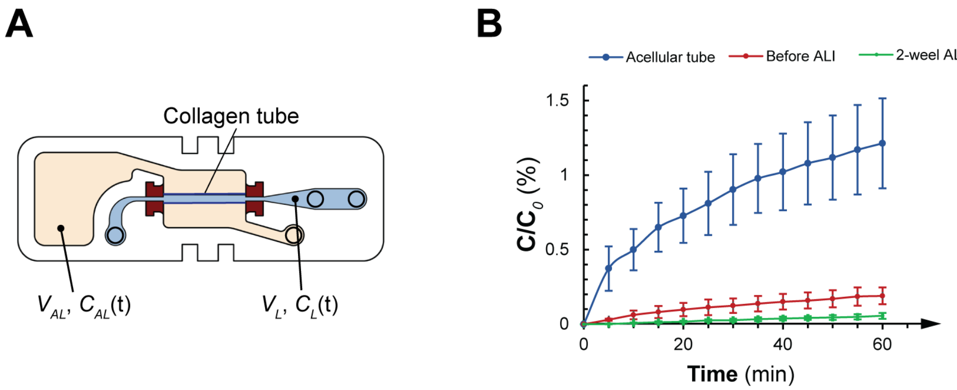


**Figure S9. Superfusion FITC-dextran concentration during the flux assay.** (A) Schematic of the dextran transport across the collagen tube wall. (B) Normalized concentration of dextran within the superfusion culture medium along the time.

### IL-8 Secretion Assay

Medium was collected from the perfusion side 5 hr after conducting the injury study with an A549 cell-based epithelium for both cases of overdistension and collapse and reopening. The collected sample was diluted 1:5 and measured using an IL-8 Human ELISA kit (# KHC0081, ThermoFisher Scientific, Waltham, MA, USA).

**YAP Localization Quantification Assay**

(1) A549 epithelium was fixed using a 4% solution of paraformaldehyde.

(2) Cells were permeabilized with 0.1 % Triton X-100 for 20 min, washed with PBS four times, then incubated in 10% goat serum for 30 min at room temperature to block nonspecific binding.

(3) Primary antibody (YAP Antibody 63.7, Santa Cruz Biotechnology, sc-101199, Dallas, TX, USA) was diluted 1:50 in PBS and incubated for 2 h at room temperature in the dark.

(4) The corresponding secondary antibody (Alexa Fluor 647 goat anti-mouse antibody, Thermo Fisher, A32728, Waltham, MA, USA) was diluted 1:200 and incubated for 1 h at room temperature in the dark.

### 16HBE14o- Immunofluorescence Staining

(1) Frozen sample sectioning & immunofluorescence staining: Tubes were fixed in 4% paraformaldehyde (PFA) for 10-20 min within the hosting device followed by a 30% sucrose solution through the lumen of the tube. Sucrose was also added to the superfusion chamber and kept overnight at 4°C. The sucrose solution was then displaced with O.C.T. Compound embedding medium (Scigen Tissue-Plus™) and transferred to a cryomold, and frozen on dry ice. 5 µm thick sections were cut using a cryostat (Leica CM1860) and mounted on slides. Citrate buffer antigen retrieval was performed, and samples were permeabilized using 0.1% Triton X-100 for 10 min. Non-specific binding was blocked using 6% normal donkey serum blocking solution for 1 h. Subsequently, a cocktail of mouse anti-pan cytokeratin (Abcam) and rabbit anti-cytokeratin 5 (Abcam) primary antibodies was diluted in the blocking solution and incubated with samples overnight at 4°C. Samples were washed twice with 1× PBS and incubated for 1 h at room temperature

with donkey anti-mouse Alexa Fluor 488 and anti-rabbit Alexa Fluor 546 secondary antibodies. Samples were washed twice with 1× PBS and counterstained with DAPI for 10 min. Samples were imaged on the EVOS M5000 Imaging System.

(2) Whole mount staining: Cells were fixed in 4% PFA within the hosting device for 10-20 min and then with 1× PBS inside the tube lumen and superfusion chamber. Cells were permeabilized with 0.1% Triton-X100 through the lumen of the tube for 10 min, followed by 6% normal donkey serum blocking solution for 1 h. Rabbit anti-ZO-1 (Proteintech) and mouse anti-Ki67 (Dako Omnis) primary antibodies diluted in blocking solution were introduced through the tube lumen and incubated overnight at 4°C. Samples were washed twice with 1× PBS and incubated for 1 h at room temperature with donkey anti-rabbit Alexa Fluor 488 and anti-mouse Alexa Fluor 546 secondary antibodies. Samples were washed twice with 1× PBS and counterstained with DAPI for 15 min. After staining was completed, the tube lumen was filled with 3% agarose and allowed to set overnight at 4°C. The tube was then excised from the hosting device and was mounted on a glass capillary for imaging on Zeiss Light Sheet Z1 Microscope (SickKids Imaging Facility). 3D rendered image created on Arivis Vision 4D (SickKids Imaging Facility).

(3) Histology Sectioning: Cells were fixed in 10% formalin within the hosting device for 10-20 minutes and then with 1× PBS inside the tube lumen and superfusion chamber. Tube lumen was filled with 3% agarose and allowed to set overnight at 4°C. Samples were embedded in paraffin and stained with hematoxylin and eosin. Sections were imaged on Nikon Upright Eclipse Microscope.
